# NUDT2 loss defines quantitative limits for dinucleoside polyphosphate action on the cGAS–STING–TBK1 axis

**DOI:** 10.64898/2026.08.18.745545

**Authors:** Paul Weiland, Renuka Dharani Shivakumar, Ekaterina Jalomo-Khayrova, Julia Schmidt, Victor Zegarra, Ying Wang, Nicole Paczia, Stephan Kiontke, Andreas Burchert, Gert Bange

## Abstract

NUDT2 is an emerging candidate for therapeutic intervention in cancer, and its inhibition or loss is known to elevate adenosine-containing dinucleoside polyphosphates (Ap_n_Ns), including diadenosine tetraphosphate (Ap_4_A). Ap_4_A is a stress- and immune-associated nucleotide metabolite proposed to act as a second messenger, raising the possibility that NUDT2 targeting may unintentionally affect important nucleotide-sensitive signaling pathways. One such pathway is cGAS-STING signaling, a central innate immune axis that detects cytosolic DNA, produces the nucleotide second messenger 2′3′-cGAMP, and drives type I interferon responses. Because cGAS-STING also contributes to antitumor immunity and is being pharmacologically targeted in cancer, we asked whether sustained Ap_4_A accumulation perturbs this pathway.

We systematically evaluated Ap_4_A and related dinucleoside polyphosphates across the cGAS-STING-TBK1 axis using biophysical, enzymatic, structural, and cellular approaches. Contrary to a previous model, STING did not bind Ap_4_A, Ap_3_A, or Ap_4_G, despite robust binding of canonical cyclic dinucleotides. Although cGAS bound these nucleotides with micromolar affinities, DNA-activated cGAMP synthesis was inhibited only at high, supra-substrate ratios. Similarly, TBK1 inhibition required extreme Ap_4_A ratios beyond physiologically relevant levels. In a THP-1 cell model, *NUDT2* knockout caused strong Ap_4_A accumulation, but the resulting intracellular dinucleoside polyphosphate levels remained below the ratios required to inhibit cGAS or TBK1 *in vitro*. This study thus distinguishes biochemical possibility from physiological relevance and argues that NUDT2-linked Ap_4_A accumulation is unlikely to directly compromise cGAS-STING pathway activity.

## Introduction

The Nudix-type hydrolase NUDT2 has emerged as a potential therapeutic target in several cancer contexts (Abu-Rahmah et al., 2024; Hidmi et al., 2023; Oka et al., 2011), and inhibition or loss of NUDT2 has been shown to cause sustained accumulation of adenosine-containing dinucleoside polyphosphates (Ap_n_Ns) (Marriott et al., 2016). Ap_n_Ns are nucleotide-based metabolites found across pro- and eukaryotic organisms, in which adenosine is connected to a second nucleoside by a polyphosphate chain, with diadenosine tetraphosphate (Ap_4_A) being the best-characterized member (Ferguson et al., 2020; Zegarra et al., 2023).

In mammalian cells, Ap_4_A accumulation has been observed in response to oxidative stress and DNA damage (Baker & Jacobson, 1986; Marriott et al., 2015; Reinalter et al., 2025), and during immune activation, exemplified by the IgE/FcεRI-triggered LysRS-Ap_4_A-HINT1-MITF axis (Lee et al., 2004; Ofir-Birin et al., 2013; Yu et al., 2019). This regulated accumulation, turnover, and pathway engagement support the proposed role of Ap_n_Ns as nucleotide-based second messengers rather than passive metabolic byproducts.

Since Ap_n_Ns structurally resemble other nucleotide substrates, cofactors, and second messengers, their accumulation could affect a broad range of nucleotide-sensitive signaling pathways. One pathway of particular relevance is cGAS-STING signaling (cyclic GMP-AMP synthase-stimulator of interferon genes), a central mammalian innate immune axis that detects cytosolic DNA as a pathogen- or damage-associated molecular pattern (PAMP/DAMP). Since the discovery of the cGAS-STING axis in 2013 (Ablasser et al., 2013; Sun et al., 2013; J. Wu et al., 2013), it has evolved from a canonical antiviral pathway into a central framework for understanding innate immunity, sterile inflammation, genome instability, and antitumor immunity (Carozza et al., 2020; Hooftman et al., 2026; Kim et al., 2020). In brief, binding of cytosolic DNA activates cGAS to synthesize the nucleotide second messenger 2′3′-cGAMP (cyclic GMP-AMP) from ATP and GTP. 2′3′-cGAMP activates STING, which recruits the protein kinase TANK-binding kinase 1 (TBK1) and thereby promotes IRF3-dependent induction of type I interferon and inflammatory gene expression. Regulation of this pathway therefore has broad implications for chronic inflammation and the shaping of the tumor microenvironment (Ablasser & Chen, 2019; Hooftman et al., 2026).

Recent work suggests that Ap_4_A directly binds STING and attenuates STING-dependent inflammatory signaling (Guerra et al., 2020), raising the possibility that Ap_4_A accumulation during NUDT2 inhibition could unintentionally modulate cGAS-STING pathway output. In addition to Ap_4_A, related Ap_n_Ns such as Ap_3_A and the mixed-base species Ap_4_G are detected in human cells and can increase under oxidative stress (Reinalter et al., 2025). Ap_4_G is of particular interest because it mirrors the ATP/GTP substrate composition of cGAS, and we therefore included it as a chemically relevant comparator.

In this study we systematically analyzed Ap_4_A, Ap_3_A, and Ap_4_G across the cGAS-STING axis by testing whether they bind or functionally influence recombinant STING, cGAS, or TBK1 using quantitative biophysical, enzymatic, and structural approaches. To assess cellular relevance, we complemented these experiments with a human NUDT2-disruption model in which Ap_n_Ns strongly accumulate intracellularly. By comparing the resulting intracellular Ap_n_N concentrations with the biochemical activity thresholds defined *in vitro*, we determined that NUDT2-linked Ap_n_N accumulation does not reach levels compatible with direct modulation of the cGAS-STING-TBK1 axis. This study therefore distinguishes biochemical possibility from physiological relevance and provides an important framework for evaluating the immunological consequences of NUDT2 inhibition.

## Results and Discussion

### Dimeric STING does not bind Ap_4_A

Because STING was proposed to be directly inhibited by Ap_4_A (Guerra et al., 2020), we first examined ligand binding to a recombinant STING panel by isothermal titration calorimetry (ITC), which directly quantifies ligand-macromolecule interactions in solution (Bastos et al., 2023). To account for species- and variant-dependent ligand recognition, and to exclude a variant-restricted Ap_n_N interaction, we purified recombinant wild-type STING from both *Homo sapiens* (*Hs*) and *Mus musculus* (*Mm*), as well as the variants *Hs* STING_R232H_ and *Mm* STING_R231A_ (Fig. 1A, B). In these variants, the conserved arginine at position 232/231 is substituted with a histidine or alanine, respectively, as these variants have previously been linked to altered cyclic-dinucleotide recognition and STING activation (Diner et al., 2013; Gao et al., 2013; Patel & Jin, 2019; Zhang et al., 2013). Because the C-terminal tail contributes to activation and autoinhibition (Yin et al., 2012; Zhao et al., 2019), we initially aimed to retain it in all cytosolic constructs. However, tail-containing human STING yielded two distinct species, consistent with partial C-terminal instability (Fig. S1.1). We therefore used truncated human STING for homogenous purification, while retaining the tail in the stable mouse constructs to test for potential tail-dependent Ap_n_N binding (Fig. 1A, B).

**Figure 1.**
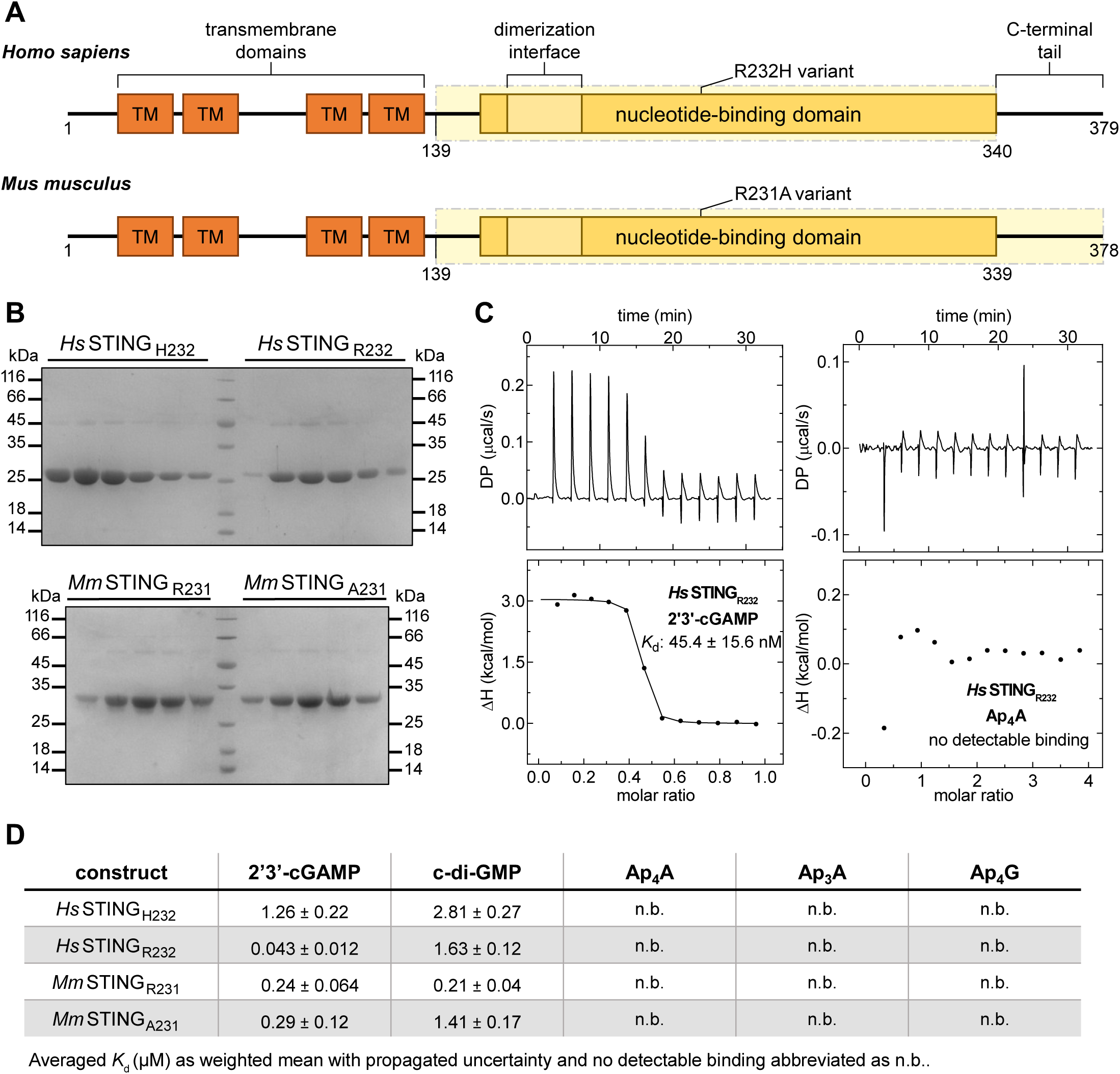
STING does not bind Ap_n_Ns. [**A**] Schematic representation of the full-length human and mouse STING, indicating domain organization and construct boundaries. The cytosolic regions used for binding analyses are highlighted in yellow, including *Hs* STING_139−340_ carrying either the wild-type R232 or variant H232 polymorphism and the corresponding *Mm* STING_139−378_ constructs carrying either R231 or A231. [**B**] SDS–PAGE analysis of purified STING constructs after size-exclusion chromatography. The proteins migrate as discrete bands at the expected molecular weights, confirming the purity of the preparations used for binding experiments. [**C**] Representative isothermal titration calorimetry (ITC) experiments of one independent replicate comparing binding of the canonical STING ligand 2′3′-cGAMP and Ap_4_A to the same human wild-type STING_R232_ construct. Raw thermograms (top) and integrated binding isotherms (bottom) demonstrate robust binding of 2′3′-cGAMP and the absence of a detectable binding signal for Ap_4_A under identical assay conditions. [**D**] Summary of ITC binding measurements for canonical ligands and dinucleoside polyphosphates across all STING constructs tested. Averaged dissociation constants (*K*_d_) of two independent replicates are shown in µM where binding was detected; n.b. indicates no detectable binding (Fig. S1.2). All measurements were performed under identical buffer conditions (20 mM HEPES, 150 mM NaCl, 10 mM MgCl_2_, pH 7.5).

We could confirm that all four STING constructs bound the canonical ligands 2′3′-cGAMP and c-di-GMP with robust enthalpic signals and apparent dissociation constants (*K*_d_) within the reported affinities for cyclic-dinucleotide interactions (Fig. 1C, D) (Burdette et al., 2011; Chin et al., 2013; Gao et al., 2013; Shu et al., 2012; Zhang et al., 2013). These experiments demonstrate that the purified proteins are ligand-competent. This finding is particularly important because none of the STING constructs showed measurable binding to the nucleotides Ap_3_A, Ap_4_A, and Ap_4_G under identical assay conditions (Figs. 1C, D; S1.2 and Table S1.1).

Previous work displayed the proposed Ap_4_A-binding mode in a monomeric representation of STING (Guerra et al., 2020), whereas mammalian STING functions as a dimer and the canonical cyclic-dinucleotide binding pocket is formed at the dimer interface (Ouyang et al., 2012; Shi et al., 2015). We therefore verified the oligomeric state of our recombinant constructs by analytical size-exclusion chromatography and found all STING variants to be dimeric in solution (Fig. S1.3). The mouse constructs eluted at a slightly higher apparent molecular weight, likely reflecting altered migration caused by their retained, disordered C-terminal tails. We then repeated the binding-mode prediction using dimeric STING in Boltz-2 (Passaro et al., 2025). The model confidently recovered the canonical 2′3′-cGAMP binding mode (mean ligand pLDDT: 93.6 ± 3.0), whereas Ap_4_A showed substantially lower ligand confidence (mean ligand pLDDT: 64.1 ± 8.3) (Fig. S1.4), further supporting the conclusion that Ap_4_A and related dinucleoside polyphosphates are unlikely to bind STING.

Together, our ITC and prediction data argue that mammalian STING does not bind Ap_4_A and related dinucleoside polyphosphates *in vitro*. Moreover, even weak or transient binding would be unlikely to compete efficiently with 2′3′-cGAMP, which is bound by STING with affinities in the nanomolar range (Fig. 1D).

### DNA-activated catalysis of cGAS is not efficiently impaired by Ap_4_A

Having found no evidence that recombinant STING binds Ap_4_A, Ap_3_A, or Ap_4_G *in vitro*, we next turned to cGAS. We first tested whether dinucleoside polyphosphates bind cGAS in the absence of DNA. Because ITC with wild-type cGAS is confounded by nucleotide turnover, we generated catalytically inactive human and mouse cGAS_QN_ mutants based on previously reported experiments (S. Wu et al., 2024). In these constructs, conserved acidic residues required for Mg^2+^ coordination and catalysis were replaced by glutamine and asparagine (E225Q/D227N in human and E211Q/D213N in mouse cGAS) (Fig. 2A, B). Under these assay conditions, the canonical substrates ATP and GTP bound human and mouse cGAS with apparent *K*_d_ values in the low-to-mid micromolar range, spanning approximately 70−170 µM (Figs. 2C, D; S2.1). Interestingly, all three dinucleosides were also bound by cGAS, with *K*_d_ values of ∼50−150 µM for human and stronger binding of ∼10−40 µM for mouse cGAS. Among these, Ap_4_G showed the highest affinity (Figs. 2C, D; S2.1, Table S2.1), potentially reflecting better accommodation of its mixed adenine-guanine composition within the ATP/GTP-binding architecture. These data show that cGAS can bind Ap_n_Ns already in the absence of DNA, consistent with recent work showing that cGAS can bind ATP and GTP nonproductively, before DNA binding organizes the active site for catalysis (Hooy & Sohn, 2018; S. Wu et al., 2024).

**Figure 2.**
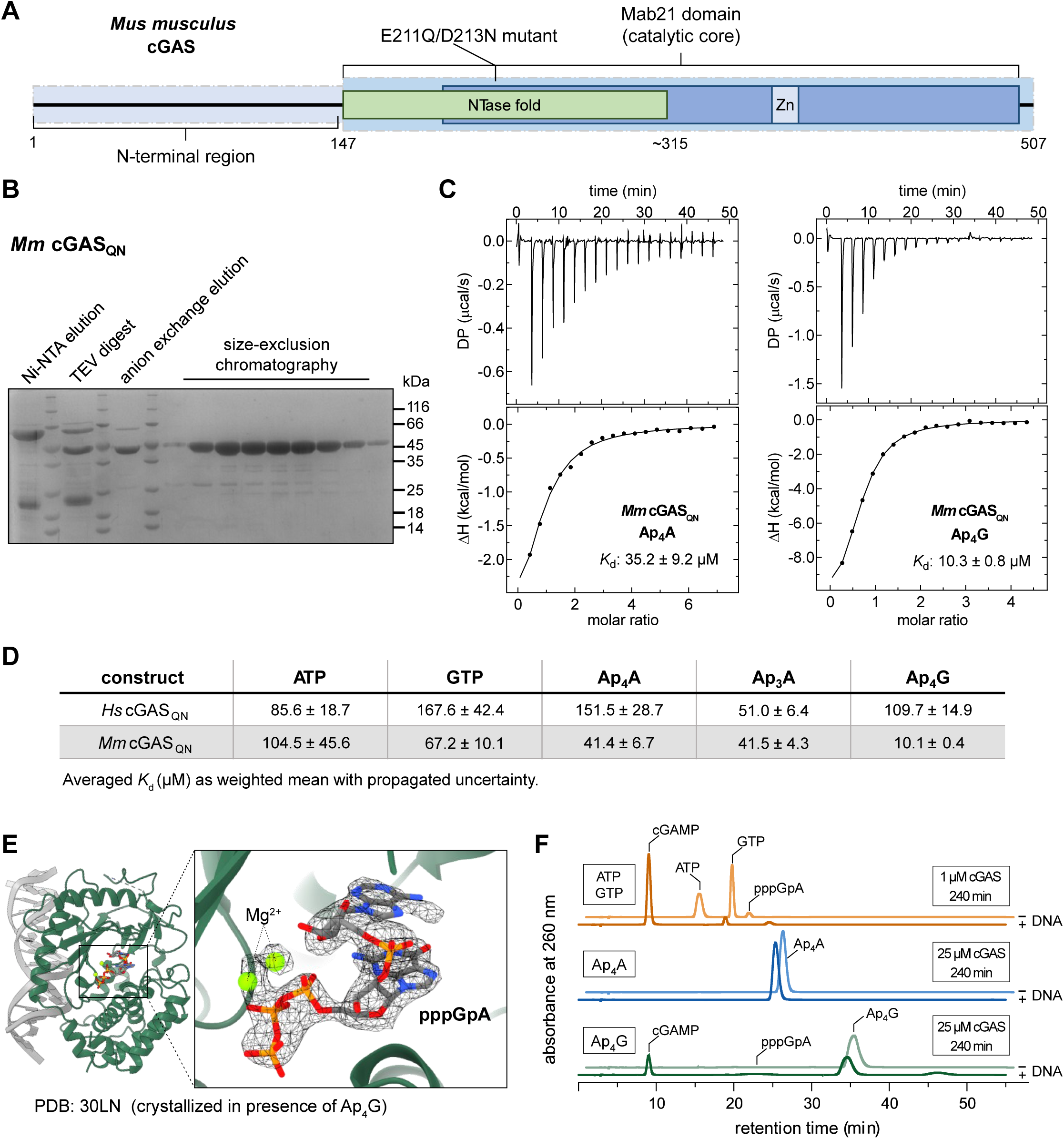
cGAS binds Ap_4_A, but its DNA-activated catalysis is not efficiently inhibited. [**A**] Schematic representation of full-length cGAS from *Mus musculus* (*Mm*) cGAS, indicating domain organization and construct boundaries. The intrinsically disordered N-terminal region (residues 1−146) is separated from the conserved Mab21 catalytic domain (residues 147−507). Within the catalytic domain, the NTase fold (∼150−320) and the C-terminal helical lobe containing the Zn-binding motif as well as the conserved acidic residues replaced by glutamine and asparagine (E211Q/D213N) are indicated. Construct boundaries used in this study are highlighted in light-blue. [**B**] Representative purification workflow for cGAS proteins used in this study, shown with an SDS–PAGE analysis of purified *Mm* cGAS_QN_. [**C**] Representative isothermal titration calorimetry (ITC) experiments showing binding of *Mm* cGAS_QN_ to Ap_4_A and Ap_4_G. [**D**] Summary of ITC binding measurements for canonical nucleotides and dinucleoside polyphosphates with human and mouse cGAS_QN_ constructs. Averaged dissociation constants (*K*_d_) of two independent replicates are shown in µM. All measurements were performed in the absence of DNA under identical buffer conditions (25 mM HEPES pH 7.5, 125 mM potassium acetate, 5 mM MgCl_2_, and 1 mM TCEP). [**E**] Structural analysis of DNA-bound wild-type cGAS crystallized in the presence of Ap_4_G. Crystallization experiments did not reveal electron density corresponding to intact Ap_4_G in the active site. Instead, density consistent with pppGpA was observed in the binding pocket with the corresponding 2*F*_o_-*F*_c_ map contoured at 1 σ shown in black. The structure was deposited in the Protein Data Bank under accession code 30LN. [**F**] HPLC analysis of cGAS reaction products generated from canonical ATP/GTP substrates or dinucleoside polyphosphates. Canonical reactions containing ATP and GTP produced cGAMP and pppGpA in the presence of DNA. Ap_4_A did not support detectable cGAMP formation, whereas Ap_4_G yielded cGAMP and pppGpA specifically in the presence of DNA. Ap_4_A and Ap_4_G reactions were performed with 25-fold higher cGAS concentration than the canonical ATP/GTP reaction to detect low-efficiency turnover.

We next sought to define the binding mode of Ap_4_A and Ap_4_G by crystallizing mouse cGAS_147−507_ (Fig. 2A) with either dinucleoside (Tab. 1). Surprisingly, in crystals obtained without DNA, the nucleotide-binding pocket remained empty, despite positive ITC binding signals in the absence of DNA. Electron density within the nucleotide binding site was only obtained in crystals of DNA-bound cGAS. However, neither structure contained a fully resolved or intact nucleotide. In the Ap_4_A dataset, the electron density corresponded only to an ATP moiety, with no detectable density for the remainder of the ligand (Fig. S2.2). In the Ap_4_G dataset, we unexpectedly found density consistent with pppGpA, the linear precursor of cGAMP (Fig. 2E). This suggests that DNA-bound cGAS may process Ap_4_G into canonical reaction intermediates and products.

**Table 1.**
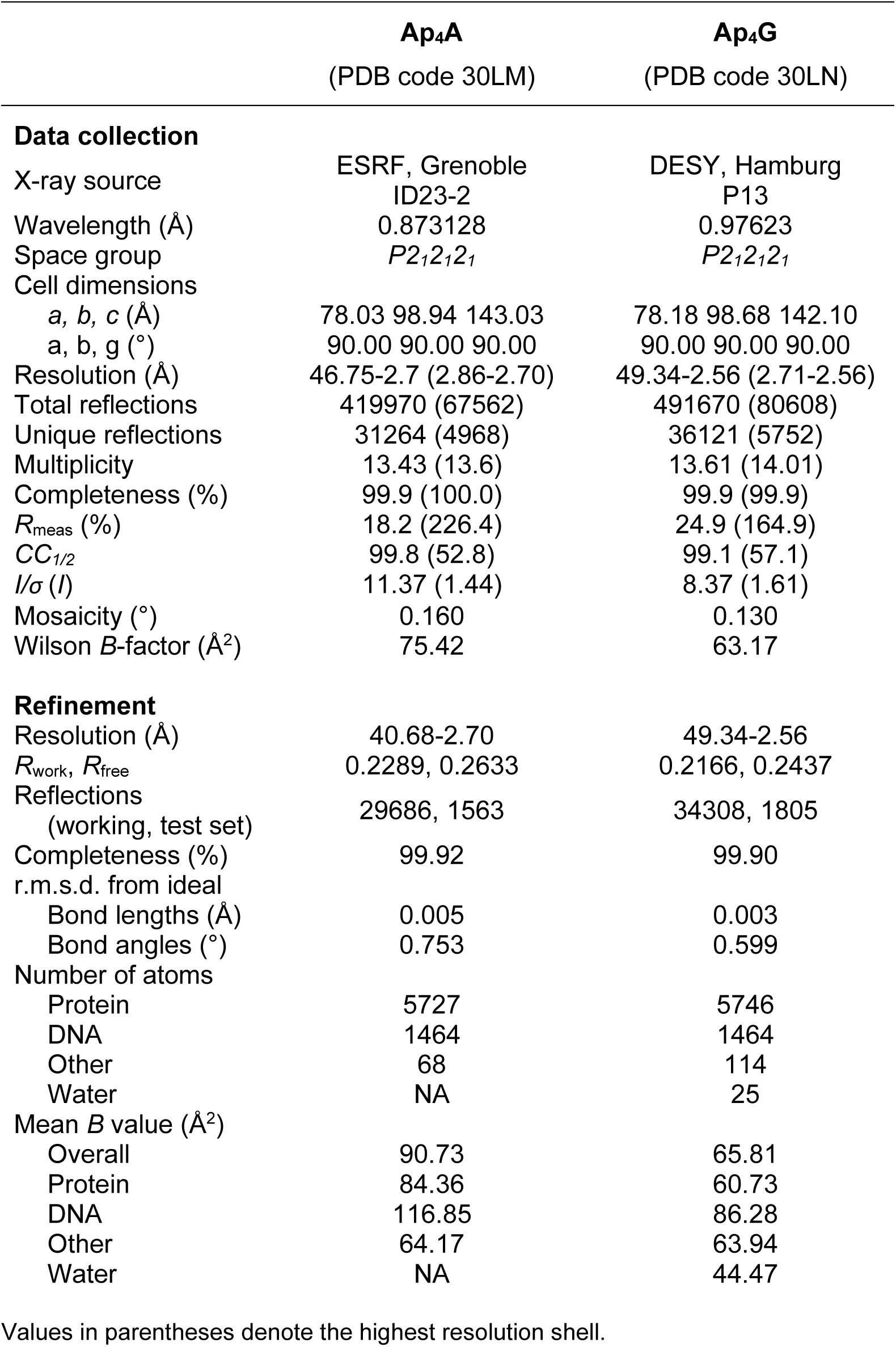
Crystallographic statistics of mouse cGAS crystallized in presence of either Ap_4_A or Ap_4_G.

We therefore tested whether Ap_4_A or Ap_4_G can serve as cGAS substrates in the absence or presence of DNA. We used the same wild-type mouse cGAS_147−507_ construct, and analyzed reaction products by High-Performance Liquid Chromatography (HPLC) (Figs. 2F; S2.3). Canonical reactions containing ATP, GTP, and DNA produced cGAMP and pppGpA as expected (Figs. 2F; S2.3). We then tested Ap_4_A and Ap_4_G at the same nucleotide concentration and over the same time course, but with 25-fold higher cGAS concentration. Under these conditions, Ap_4_A did not support cGAMP synthesis and remained entirely intact (Figs. 2F; S2.3) while Ap_4_G yielded detectable cGAMP as well as pppGpA specifically in the presence of DNA (Figs. 2F; S2.3). However, product formation was far lower than in canonical ATP/GTP reactions despite the increased enzyme concentration. Thus, cGAS binds Ap_4_A, but it is not a productive substrate, whereas Ap_4_G is not only bound by cGAS but also inefficiently processed when supplied at high concentration, consistent with the pppGpA density observed in the crystal structure.

This inefficient, substrate-like behavior prompted us to ask how these dinucleosides influence the canonical ATP/GTP-driven cGAS catalysis. We used full-length wild-type mouse cGAS (Fig. 2A) supplied with ATP, GTP, and DNA, and monitored activity using a pyrophosphatase-coupled malachite green assay. Because cGAS and TBK1 operate in the presence of abundant substrate pools, absolute Ap_n_N concentration alone does not define inhibitory relevance. We therefore expressed inhibitory potency as D/S_50_, the dinucleoside:substrate ratio required for 50 % inhibition, allowing direct comparison with intracellular nucleotide ratios. Ap_4_A and Ap_3_A reduced cGAS activity only when present in large excess over the canonical substrates, with D/S_50_ values of ∼46 and ∼27, respectively (Fig. 3A). Curve fitting was robust for Ap_4_A and Ap_4_G, with R^2^ values of 0.95 and 0.97, respectively, whereas the weaker and less complete inhibition by Ap_3_A resulted in a poorer fit with an R^2^ of 0.67. Notably, Ap_4_G inhibited more effectively, with a D/S_50_ of ∼6, consistent with its stronger apparent binding affinity (Figs. 2D and 3A).

**Figure 3.**
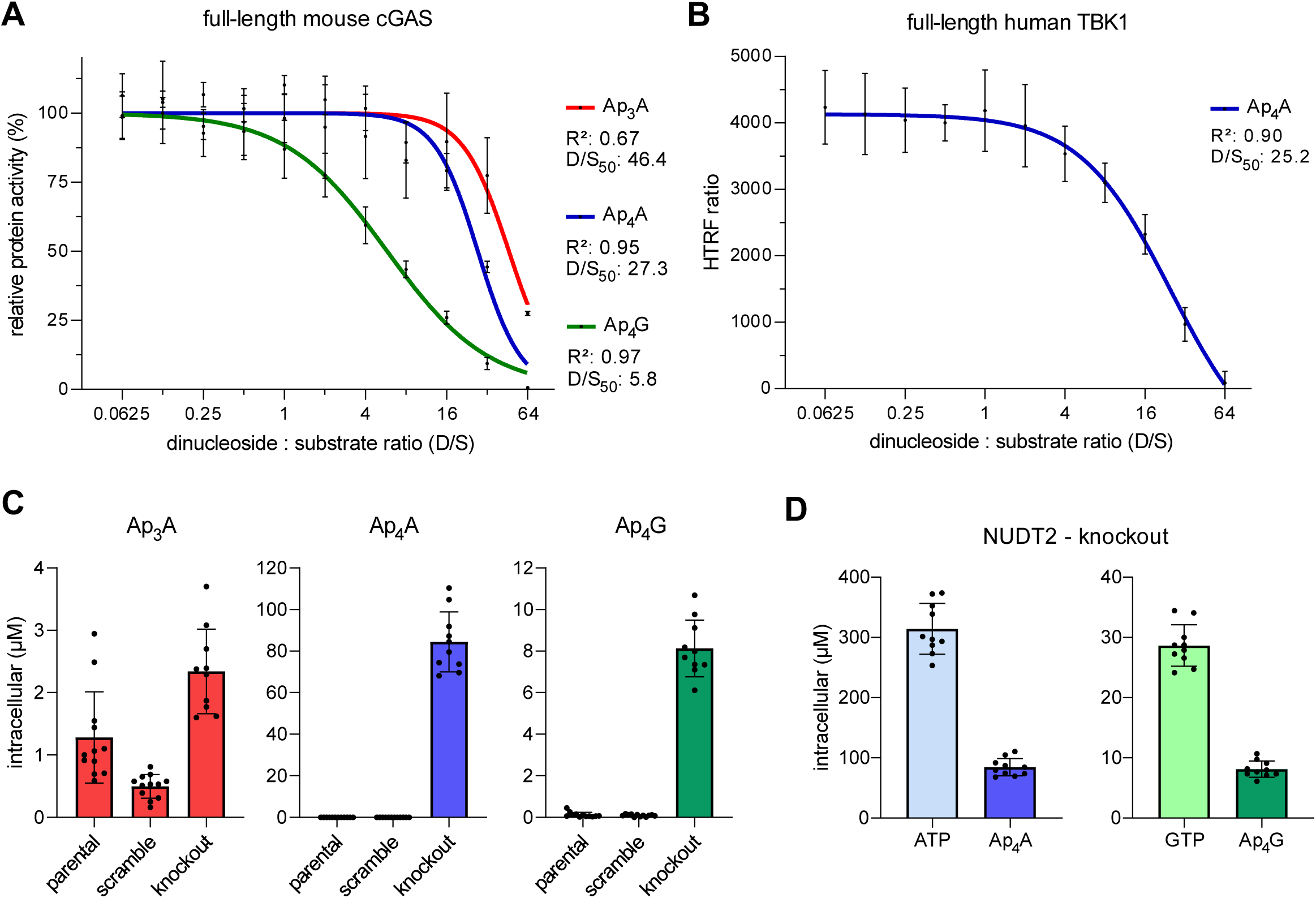
Intracellular Ap_4_A and Ap_4_G concentrations remain below cGAS and TBK1 inhibitory thresholds. [**A**] Inhibition of DNA-activated full-length mouse cGAS by Ap_3_A, Ap_4_A, and Ap_4_G, measured using a pyrophosphatase-coupled malachite green assay. Normalized relative activity is based on absorbance measurements and was plotted against the dinucleoside:substrate ratio (D/S). D/S_50_ values indicate the molar dinucleoside:substrate ratio required to reduce cGAS activity by 50 %. Curves were fitted by least-squares nonlinear regression using a variable-slope inhibitor-response model, with D/S used as the inhibitor variable and R^2^ values indicate goodness of fit. Error bars represent means with standard deviations from three independent experiments, each measured in duplicate. [**B**] Inhibition of full-length human TBK1 by Ap_4_A, measured using a Homogeneous Time-Resolved Fluorescence (HTRF) kinase assay. The HTRF ratio is plotted against the Ap_4_A:ATP ratio, with the D/S_50_ indicating the ratio required for 50 % inhibition. The curve was fitted by least-squares nonlinear regression using a variable-slope inhibitor-response model, with D/S used as the inhibitor variable and R^2^ values indicate goodness of fit. Error bars represent means with standard deviations from three independent experiments, each measured in triplicate. [**C**] LC-QQQ-MS quantification of intracellular Ap_3_A, Ap_4_A, and Ap_4_G concentrations in parental, scrambled sgRNA control, and NUDT2-knockout THP-1 cells. NUDT2 loss strongly elevates Ap_4_A and Ap_4_G, whereas Ap_3_A levels remain unchanged. [**D**] Comparison of intracellular ATP with Ap_4_A and GTP with Ap_4_G in NUDT2-knockout THP-1 cells. Both dinucleosides reach approximately one-third of the corresponding nucleotide pools, remaining well below the inhibitory ratios required for cGAS or TBK1 inhibition *in vitro*. [**C**+**D**] Parental and scrambled sgRNA control cells were represented by six independent single-cell clones and NUDT2-knockout cells by five; each clone was extracted in duplicate. Bars represent means with standard deviations and individual points represent duplicate extracts from independent single-cell clones.

Together, these data demonstrate that cGAS binds Ap_n_Ns and, in the case of Ap_4_G, can process them inefficiently under high-concentration conditions. However, inhibition of canonical cGAMP synthesis requires Ap_n_Ns to be present in excess over the ATP and GTP substrates, arguing against potent inhibition of cGAS under physiological nucleotide ratios and concentrations (Lane & Fan, 2015; Traut, 1994).

### Activity of TBK1 is only inhibited at extreme Ap_4_A excess

TBK1 is the central kinase downstream of STING and uses ATP as phosphate donor, making it a plausible target for Ap_4_A-dependent nucleotide interference. We therefore examined whether Ap_4_A directly affects TBK1 activity using full-length human TBK1 in a Homogeneous Time-Resolved Fluorescence (HTRF) kinase assay. Under conditions supporting ATP-dependent substrate phosphorylation, Ap_4_A reduced TBK1 activity only when present in strong excess over ATP, with a D/S_50_ value of ∼25 (Fig. 3B).

This steep ratio requirement *in vitro* argues against selective high-affinity regulation of TBK1 by Ap_4_A as the required ratios are far above physiologically relevant nucleotide conditions (Lane & Fan, 2015; Traut, 1994).

### NUDT2 loss elevates Ap_4_A and Ap_4_G but remains below cGAS and TBK1 inhibitory thresholds

To assess whether NUDT2 loss elevates Ap_n_Ns to ratios capable of inhibiting cGAS or TBK1 *in vitro*, we next generated a CRISPR/Cas9 THP-1 NUDT2 knockout model. Loss of NUDT2 was confirmed by both western blotting and the expected accumulation of NUDT2-linked Ap_n_Ns (Figs. 3C, D and S3.1). Targeted nucleotide analysis showed that NUDT2 loss strongly elevated intracellular Ap_4_A, and Ap_4_G concentrations, whereas these dinucleosides were low or undetectable in parental and control cells (scrambled gRNA) (Fig. 3D). Based on monitored cell diameters and calculated cell volume, Ap_4_A accumulated to apparent intracellular concentrations in the range of 70−100 µM, whereas Ap_4_G only accumulated to a low micromolar range of 7−10 µM (Fig. 3D). The Ap_4_A concentrations fall within the micromolar range reported previously for mammalian cells (Lee et al., 2004; Marriott et al., 2015, 2016). Ap_3_A levels were largely unchanged in the knockout and remained in the lower single-digit micromolar range (Fig. 3D), consistent with the Ap_4_N-type substrate preference of NUDT2 (Carreras-Puigvert et al., 2017; Guranowski et al., 2000). Interestingly, Ap_4_A and Ap_4_G accumulation mirrored the relative availability of ATP and GTP (Fig. 3E). The lower abundance of GTP relative to ATP was paralleled by lower Ap_4_G accumulation relative to Ap_4_A, consistent with substrate-pool availability contributing to Ap_n_N accumulation.

Despite strong Ap_4_A and Ap_4_G accumulation, the relevant intracellular dinucleoside:substrate ratios remained below the inhibitory thresholds measured biochemically (Fig. 3). ATP and GTP levels were not substantially altered between the three cell lines (Fig. S3.2), and in NUDT2 knockout cells Ap_4_A and Ap_4_G reached only approximately one-third of the corresponding ATP and GTP pools, respectively (Fig. 3E). These ratios are far below the D/S_50_ values required for cGAS inhibition by Ap_4_A/Ap_4_G or TBK1 inhibition by Ap_4_A *in vitro* (Fig. 3A, B).

Together with the absence of detectable binding of recombinant STING to Ap_n_Ns (Figs. 1C, D and S.1.2), these data argue that NUDT2-linked Ap_n_N accumulation is unlikely to directly suppress the cGAS-STING-TBK1 axis through STING binding or cGAS/TBK1 inhibition. Our study therefore defines quantitative limits for Ap_n_N action and suggests that therapeutic NUDT2 inhibition is unlikely to unintentionally impair this pathway through direct inhibition of STING, cGAS, or TBK1.

## Material and Methods

### Cloning of recombinant STING and cGAS constructs

Human STING (UniProt: Q86WV6; residues 139−379) was amplified from synthetic DNA obtained from Integrated DNA Technologies (IDT), whereas mouse STING (UniProt: Q3TBT3; residues 139−378) was amplified from cDNA. All STING constructs were cloned into pET24d-based expression vectors encoding an N-terminal 6xHis tag using a BsaI-based scarless Golden Gate cloning strategy (see below).

Human cGAS (UniProt: Q8N884; residues 157−522) was amplified from cDNA, whereas mouse cGAS (UniProt: Q8C6L5; residues 2−507) was amplified from a synthetic DNA fragment obtained from IDT, with partial codon optimization of the N-terminal coding region. The individual cGAS constructs were cloned into modified expression vectors encoding an N-terminal 6xHis-SUMO tag followed by a Tobacco Etch Virus (TEV) protease cleavage site. The modified expression vectors and scarless BsaI-based Golden Gate cloning strategy are described in more detail elsewhere (Zweng et al., 2025). Truncated constructs and point-mutated variants were subsequently amplified from these recombinant plasmids and assembled into the corresponding expression vectors by the same BsaI-dependent Golden Gate strategy.

PCR amplification was performed with Q5 High-Fidelity Master Mix (New England Biolabs) using construct-specific primer pairs and annealing temperatures of 60−64°C. Point mutations, including human and mouse STING allelic variants and cGAS catalytic-site variants, were introduced using overlapping mutagenic primers. For each mutation, two overlapping PCR fragments were amplified and assembled into the corresponding expression vector by BsaI-dependent Golden Gate cloning. PCR products and plasmids were purified using GeneJET purification kits (Thermo Fisher Scientific). Cloning was performed in chemically competent *Escherichia coli* DH5α cells (New England Biolabs), and all final constructs were verified by Sanger sequencing (Microsynth). Recombinant plasmids used in this study are listed in Table 2.

**Table 2.**
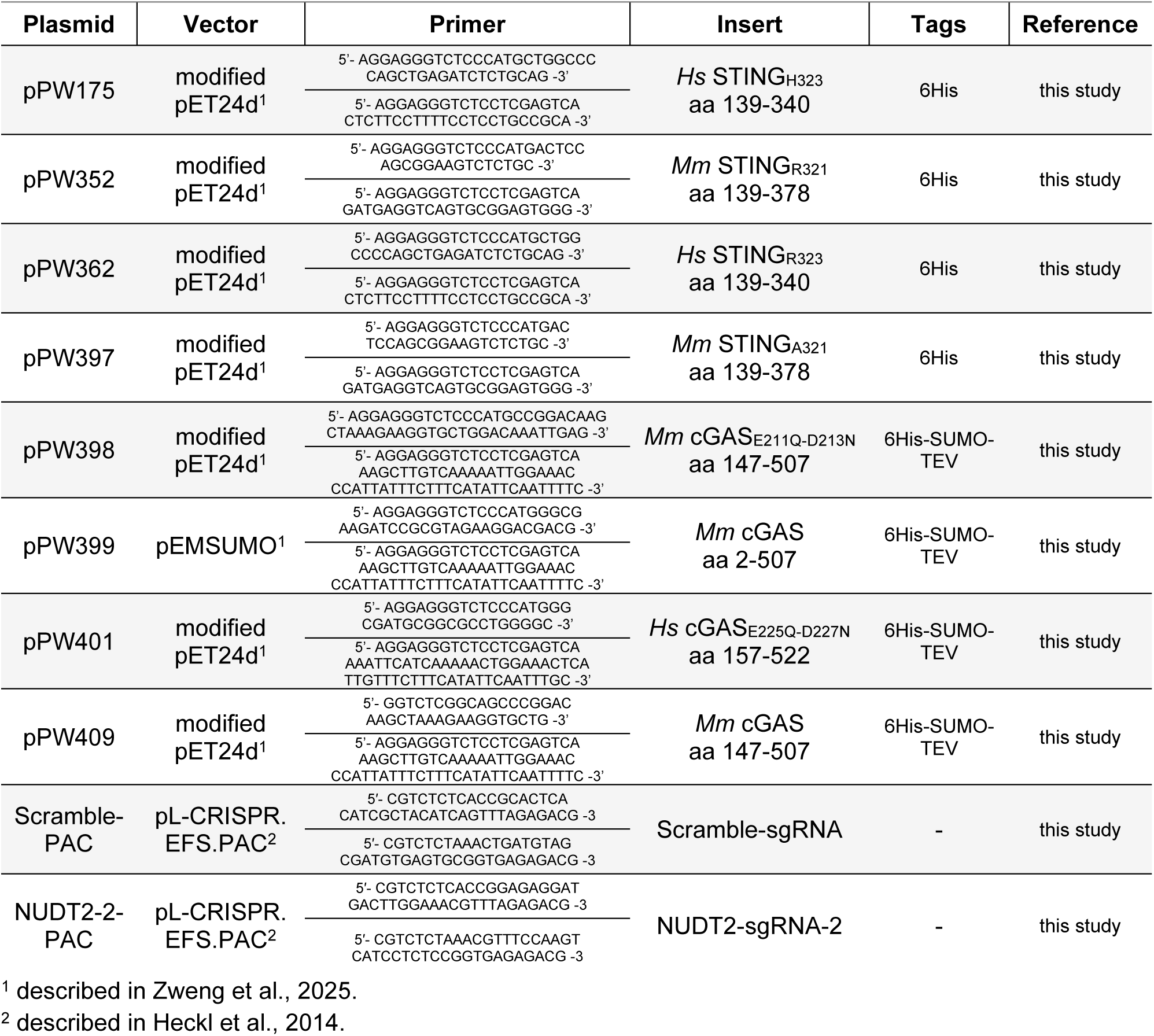
Plasmids used in this study. The table below lists all plasmids and their corresponding primer pairs.

No experiments involving live animals or human participants were performed in this study. Human and mouse designations refer to recombinant protein sequence origin, synthetic DNA or cDNA templates.

### Expression and purification of recombinant STING proteins

For recombinant protein expression, plasmids encoding the respective STING constructs were transformed into chemically competent *E. coli* BL21(DE3) cells (Promega). Transformants were selected on LB agar (Luria/Miller, Roth) supplemented with kanamycin (50 µg/mL). For each construct, 2 L of LB medium (10 g/L tryptone, 5 g/L yeast extract, and 10 g/L NaCl) supplemented with kanamycin (50 mg/L) and 10 g/L α-lactose were inoculated directly from colonies grown on selection plates. Cultures were grown overnight at 30°C with shaking to allow autoinduction of protein expression.

Cells were harvested by centrifugation at 4,500 x *g* for 15 min at 4°C and resuspended in Buffer A (20 mM HEPES, 500 mM NaCl, 5 % (v/v) glycerol, 40 mM imidazole pH 8.0). Cell lysis was performed using an LM10 Microfluidizer (Microfluidics) at 15,000 psi. Lysates were clarified by centrifugation at 47,850 x *g* for 20 min at 4°C and the cleared lysate was applied to two 1 mL HisTrap FF columns (Cytiva). After washing with five column volumes of Buffer A, bound protein was eluted using 20 mL Buffer B (20 mM HEPES, 250 mM NaCl, 250 mM imidazole pH 8).

The eluted protein was subsequently concentrated using Amicon Ultra-10K centrifugal concentrators (Millipore) and further purified by size-exclusion chromatography on a HiLoad 16/600 Superdex 200 pg column (Cytiva) equilibrated in 20 mM HEPES, 150 mM NaCl, and 10 mM MgCl_2_ pH 7.5. Peak fractions containing purified STING were pooled, concentrated, flash-frozen in liquid nitrogen, and stored at −80°C until further use. Protein concentrations were determined spectrophotometrically using a NanoDrop Lite instrument (Thermo Fisher Scientific).

### Expression and purification of recombinant cGAS proteins

For recombinant protein expression, plasmids encoding the respective cGAS constructs were transformed into chemically competent *E. coli* Rosetta pLysS cells (Novagen). Transformants were selected on LB agar (Luria/Miller, Roth) supplemented with ampicillin (100 µg/mL) and chloramphenicol (34 µg/mL). Colonies were used to inoculate overnight cultures in LB medium (10 g/L tryptone, 5 g/L yeast extract, and 10 g/L NaCl) shaking at 37°C. For each expression, 6−12 L of LB medium supplemented with ampicillin and chloramphenicol were inoculated from the overnight culture to an initial OD_600_ of 0.05 and grown at 37°C with shaking until an OD_600_ of 0.6−0.8 was reached. Cultures were cooled to 18°C, protein expression was induced with 0.5 mM IPTG, and cells were cultured for an additional 16 h at 18°C.

Cells were harvested by centrifugation at 4,500 x *g* for 15 min at 4°C and resuspended in Buffer A (20 mM Tris-HCl pH 8.0, 500 mM NaCl, 5 % (v/v) glycerol, 40 mM imidazole, 1 mM TCEP). Cell lysis was performed using an LM10 Microfluidizer (Microfluidics) at 15,000 psi. Lysates were clarified by centrifugation at 47,850 x *g* for 20 min at 4°C and applied to four 1 mL HisTrap FF columns (Cytiva). After washing with five column volumes of Buffer A, bound protein was eluted using Buffer B (20 mM Tris-HCl pH 8.0, 250 mM NaCl, 250 mM imidazole, 1 mM TCEP). The eluate was incubated with recombinant TEV protease (20 µg/mL) for 3−4 h at room temperature to remove the N-terminal 6xHis-SUMO tag. The digestion mixture was subsequently buffer-exchanged into Buffer C (20 mM Tris-HCl pH 7.5, 150 mM NaCl, and 1 mM TCEP) using Amicon Ultra-30K centrifugal concentrators (Millipore) and further purified by anion-exchange chromatography on two 5 mL HiTrap Q FF columns (Cytiva) using an NGC chromatography system (Bio-Rad). Flow-through and wash fractions containing the cleaved tag and other contaminants were discarded. Bound protein was eluted by increasing the concentration of Buffer D (20 mM Tris-HCl pH 7.5, 1 M NaCl, and 1 mM TCEP). For the mouse cGAS construct used for crystallization, anion-exchange chromatography was replaced by purification using two 1mL HiTrap Heparin HP columns (Cytiva) using the same buffer system and NaCl gradient. Peak fractions containing cGAS were pooled, concentrated, and subjected to size-exclusion chromatography (SEC) on a HiLoad 16/600 Superdex 200 pg column (Cytiva) using construct specific buffers.

Human cGAS (E225Q-D227N, residues 157−522) and mouse cGAS (E211Q-D213N, residues 147−507) variants used for ITC are based on the work of Jungsan Sohn (Johns Hopkins) (S. Wu et al., 2024) and were purified in Size-Exclusion Chromatography (SEC) buffer containing 25 mM HEPES pH 7.5, 125 mM potassium acetate, 5 mM MgCl_2_, and 1 mM TCEP. Full-length mouse cGAS (residues 2−507) used for activity assays was purified in buffer containing 25 mM HEPES pH 7.5, 150 mM NaCl, 10 mM MgCl_2_, and 1 mM TCEP. Mouse cGAS (residues 147−507) used for crystallization was purified in buffer containing 20 mM Tris-HCl pH 7.5, 150 mM NaCl, 5 mM MgCl_2_, and 0.5 mM TCEP.

Fractions containing purified cGAS were pooled, concentrated using Amicon Ultra-30K centrifugal concentrators (Cytiva), flash-frozen in liquid nitrogen, and stored at −80°C until further use. Protein concentrations were determined spectrophotometrically using a NanoDrop Lite instrument (Thermo Fisher Scientific).

### Analytical size-exclusion chromatography of STING constructs

The oligomeric state of purified STING constructs was analyzed by analytical size-exclusion chromatography using an ÄKTA pure™ (Cytiva) chromatography system equipped with a Superdex 75 Increase 10/300 GL column (Cytiva). The column was equilibrated in 20 mM HEPES pH 7.5 and 150 mM NaCl.

The column was calibrated using globular protein standards of known molecular weight: cytochrome c (12.4 kDa, Sigma-Aldrich), carbonic anhydrase (29 kDa, Cytiva), ovalbumin (43 kDa, Cytiva), conalbumin (75 kDa, Cytiva) and aldolase (158 kDa, Cytiva). Elution peak maxima were determined from the chromatograms and analyzed using GraphPad Prism. A standard curve was generated by plotting the logarithm of the molecular weight against the corresponding elution volume of the calibration standards. Apparent molecular weights of the STING constructs were calculated from this calibration curve and compared with the theoretical monomeric and dimeric molecular weights to infer their apparent oligomeric state in solution.

### Isothermal titration calorimetry (ITC)

The thermodynamic parameters of ligand binding to STING and cGAS constructs were determined using a MicroCal PEAQ-ITC instrument (Malvern Panalytical). For STING-binding measurements, *Hs* STING_H232_, *Hs* STING_R232_, *Mm* STING_R231_, and *Mm* STING_A231_ were each used at a protein concentration of 50 µM. The ligands 2′3′-cGAMP, c-di-GMP, Ap_3_A, Ap_4_A, and Ap_4_G (Jena Bioscience) were used at concentrations ranging from 250 µM to 1 mM. Proteins and ligands were prepared in filtered (0.22 µm) size-exclusion chromatography buffer containing 20 mM HEPES pH 7.5, 200 mM NaCl, 10 mM MgCl_2_. Following an initial 0.4 µL pre-injection, 12 injections of 3 µL were performed at 150s intervals. For cGAS-binding measurements, the previously described catalytically inactive *Hs* cGAS_QN_ and *Mm* cGAS_QN_ variants were used (Fig. 2A, B) (S. Wu et al., 2024). *Hs* cGAS_QN_ was used at concentrations ranging from 50 to 80 µM, whereas *Mm* cGAS_QN_ was used at 50 µM. ATP, GTP, Ap_3_A, Ap_4_A, and Ap_4_G were used at concentrations ranging from 1.2 to 3.5 mM. Proteins and ligands were prepared in filtered (0.22 µm) size-exclusion chromatography buffer containing 25 mM HEPES pH 7.5, 125 mM potassium acetate, 5 mM MgCl_2_, and 1 mM TCEP. Following an initial 0.4 µL pre-injection, 12−18 injections of 3 or 2 µL, respectively, were performed at 150s intervals.

Experiments were conducted at 25 °C with a stirring speed of 750 rpm and for all measurements, protein and ligand solutions were loaded into the sample cell and injection syringe, respectively. Data were analyzed using MicroCal PEAQ-ITC Analysis Software version 1.41 (Malvern Panalytical) with a one-set-of-sites binding model and plotted using GraphPad Prism.

The ITC assays were performed from two replicates, and the resulting thermodynamic values are presented in tables S1.1 and S2.1.

### Modeling of nucleotide-bound states of cytosolic STING using Boltz-2

Nucleotide-bound states of cytosolic STING were modeled using Boltz-2 via the Neurosnap online server.(Mirdita et al., 2022; Neurosnap, 2022; Passaro et al., 2025) Human STING residues 139−340 were provided as two identical chains to model the STING dimer. Ap_4_A and 2′3′-cGAMP were supplied as small molecules using SMILES strings derived from PubChem CID 21706 and CID 135564529, respectively. Separate predictions were performed for each ligand using identical protein sequences and parameters. MSAs were generated using mmseqs2_uniref_env. No covalent restraints, pocket restraints, cyclic biopolymers, residue modifications, custom MSAs, or inference-time potentials were used. Predictions were performed with five diffusion samples, five affinity diffusion samples, six recycles, 200 sampling steps, 200 affinity sampling steps, and a step scale of 1.5. Models were evaluated using server-reported confidence metrics and inspected for ligand placement within the STING dimer interface.

### cGAS activity assay using pyrophosphatase-coupled malachite green detection

DNA-activated cGAS activity in presence of dinucleoside polyphosphates (Ap_3_A, Ap_4_A, or Ap_4_G, Jena Bioscience) was measured using a pyrophosphatase-coupled malachite green assay. In this assay, pyrophosphate released during cGAS-mediated nucleotide condensation is hydrolyzed by pyrophosphatase, and the resulting inorganic phosphate is detected using malachite green. Reactions were performed in assay buffer containing 25 mM HEPES pH 7.5, 150 mM NaCl, 5 mM MgCl_2_, and 1 mM TCEP. Full-length wild-type mouse cGAS and 60-bp dsDNA were used at final concentrations of 200 nM each. ATP and GTP were supplied at final concentrations of 100 µM each. The 60-bp dsDNA sequence used is: 5’- ATG GAA GAT CCG CGT AGA AGG ACG ACG GCG CCA CGC GCT AAG AAG CCG TCC GCG AAG CGC -3’ (S. Wu et al., 2024).

Ap_3_A, Ap_4_A, or Ap_4_G was serially diluted twofold starting from 8.54 mM, in assay buffer containing 133.34 µM ATP and 133.34 µM GTP. A condition lacking dinucleoside polyphosphates was included as the 0 mM control. Additionally, a condition lacking dinucleoside polyphosphates was stopped immediately with 50 mM EDTA and used as a negative turnover control. Reactions were assembled and initiated by mixing 15 µL of the nucleotide/dinucleoside polyphosphate dilution with 5 µL enzyme mix containing full-length mouse cGAS, 60-bp dsDNA, and yeast pyrophosphatase (Thermo Scientific) in assay buffer. Final reactions had a volume of 20 µL and contained 200 nM cGAS, 200 nM dsDNA, 100 µM ATP, 100 µM GTP, 0.05 units of pyrophosphatase per reaction, and dinucleoside concentrations ranging from 6.4 to 0 mM.

Reactions were incubated for 60 min at 37 °C, and stopped by adding 5 µL 250 mM EDTA. Stopped reactions were frozen at −80 °C until malachite green analysis.

The malachite green detection reagent was prepared by mixing 0.045 % (w/v) malachite green oxalate in ddH_2_O, with 4.2 % (w/v) ammonium molybdate in 34.5 % (v/v) concentrated HCl, and 4 % (v/v) Tween-20 in ddH_2_O. Ammonium molybdate and malachite green oxalate solutions were mixed at a 1:3 volume ratio and stirred for 20 min at room temperature. Tween-20 solution was then added at 0.1 volumes relative to the ammonium molybdate/malachite green mixture. The working detection reagent was protected from light and stored at 4 °C until use.

For phosphate detection, stopped reactions were thawed at room temperature and briefly mixed. The complete 25 µL stopped reaction was transferred to a 96-well plate containing 200 µL malachite green working detection reagent. After 10 min at room temperature, 25 µL 34 % (w/v) trisodium citrate dihydrate in ddH_2_O was added and further incubated for 45 min at room temperature, then absorbance was measured at 620 nm.

For each independent serial dilution experiment, absorbance values were first background-corrected by subtracting the negative turnover control. Activity was then normalized to the corresponding reaction lacking dinucleoside polyphosphate, which was defined as 100 % activity. Dose–response curves were fitted in GraphPad Prism using nonlinear regression with a four-parameter variable-slope inhibitor-response model. Fit was obtained by least-squares regression, and goodness of fit is reported as R^2^. D/S_50_ values denote the molar dinucleoside:substrate ratio required to reduce cGAS activity by 50 %.

### Crystallization and X-ray diffraction of cGAS complexes

Crystallization of DNA-bound mouse cGAS in presence of either Ap_4_A or Ap_4_G (Jena Bioscience) was performed by hanging-drop vapor diffusion at 20°C in 1−2 µL drops, consisting of protein complex and reservoir solution mixed in a 1:1 and 1:2 ratio. The cGAS protein used for crystallization corresponded to *Mus musculus* cGAS (UniProt Q8C6L5; residues 147−507) and included an N-terminal SGS linker remaining after TEV cleavage. For both crystallization setups, cGAS was preincubated with an 18-bp dsDNA oligonucleotide with the sequence 5′-ATC TGT ACA TGT ACA GAT-3′ (S. Wu et al., 2024) in buffer containing 20 mM Tris-HCl pH 7.5, 150 mM NaCl, 0.5 mM TCEP, and 5 mM MgCl_2_. The protein complex solution contained 250 µM cGAS, 300 µM dsDNA, and 2 mM Ap_4_A or Ap_4_G.

Crystals of the cGAS-DNA complex obtained in the presence of Ap_4_A were grown in 50 mM calcium acetate, 100 mM imidazole pH 8.0, and 30−35 % (v/v) 2-ethoxyethanol. Diffraction data for these crystals were collected under cryogenic conditions at beamline ID23-2, operated by the European Synchrotron Radiation Facility (ESRF), Grenoble, France. In this structure, ATP was modeled into the active-site pocket (PDB: 30LM).

Crystals of the cGAS-DNA complex obtained in the presence of Ap_4_G were grown in 0.1 M HEPES pH 6.5 and 30−35 % (v/v) MPD. Diffraction data for these crystals were collected under cryogenic conditions at beamline P13, operated by EMBL Hamburg at PETRA III, DESY. In the resulting structure, electron density in the active-site region was consistent with pppGpA and was therefore modeled into the binding pocket (PDB: 30LN). Ligand restraints for pppGpA were generated with Grade2 (Smart et al., 2021) from a SMILES string, and a new corresponding PDB ligand code was generated for deposition.

X-ray diffraction data were processed with XDS and XSCALE (Kabsch, 2010a, 2010b). Initial phases were determined by molecular replacement using PHASER (McCoy et al., 2007)with the mouse cGAS-DNA structure PDB 7UTT as search template. The final structures were obtained through iterative model building in COOT (Emsley et al., 2010) and refinement using PHENIX (Liebschner et al., 2019). The final structures were deposited in the RCSB Protein Data Bank under accession codes 30LM and 30LN. Structural figures were rendered and visualized with UCSF ChimeraX (Pettersen et al., 2021).

### HPLC analysis of substrate specificity by cGAS

To test whether Ap_4_A or Ap_4_G (Jena Bioscience) can serve as substrates for cGAS, reactions were performed using mouse cGAS_147−507_, the same construct used for crystallization. Reactions were carried out in containing 25 mM HEPES pH 7.5, 150 mM NaCl, 5 mM MgCl_2_, and 1 mM TCEP.

Reactions were assembled from separate protein and nucleotide/DNA mixtures and initiated by mixing equal volumes at 37 °C. For control reactions with canonical substrates, reactions contained 1 µM cGAS_147−507_, 1 µM 60-bp dsDNA, 1 mM ATP, and 1 mM GTP. For reactions testing Ap_4_A or Ap_4_G as potential substrates, reactions contained 25 µM cGAS_147−507_, 25 µM dsDNA, and 1 mM Ap_4_A or Ap_4_G. Matching reactions lacking DNA were prepared by replacing dsDNA with ddH_2_O. The 60-bp dsDNA sequence used is: 5’- ATG GAA GAT CCG CGT AGA AGG ACG ACG GCG CCA CGC GCT AAG AAG CCG TCC GCG AAG CGC -3’ (S. Wu et al., 2024).

Reactions were incubated at 37 °C, and time points were sampled at 0, 15, 30, 60, 120, and 240 min. At each time point, 10 µL reaction mixture was quenched by mixing with 20 µL acetonitrile. Samples were further diluted with 20 µL ddH_2_O and centrifuged at 16,500 x *g* for 5 min. Supernatants were carefully transferred to High-Performance Liquid Chromatography (HPLC) vials (Macherey-Nagel) for analysis.

Reaction products were analyzed by HPLC using an Agilent 1260 Infinity system equipped with a Metrosep A Supp 5-150/4.0 column and a 10-µL injection volume. To optimize nucleotide retention and improve the chromatographic resolution of the reaction products, a gradient method was applied at a flow rate of 0.6 mL/min using 90 mM ammonium carbonate (pH 9.25; mobile phase A) and 56 mM ammonium carbonate (pH 9.25; mobile phase B). The column was initially equilibrated with 100% mobile phase A, which was maintained for 13 min. The mobile phase was changed to 100% B from 13.0 to 13.1 min and maintained until 50 min. The column was then returned to 100% A from 50.0 to 50.1 min and re-equilibrated until 55 min. Chromatograms were recorded by absorbance at 260 nm and pure nucleotide standards were used for peak identification. All nucleotides were bought from Jena Bioscience except pppGpA which was bought from BioLog.

### TBK1 kinase activity assay

TBK1 kinase activity was measured using the Homogeneous Time-Resolved Fluorescence (HTRF) KinEASE-STK S1 kit (Revvity) according to the manufacturer’s protocol, with modifications to test Ap_4_A-dependent inhibition. The assay is based on phosphorylation of the biotinylated STK substrate, followed by detection with STK antibody-cryptate and streptavidin-XL665. The resulting time-resolved FRET signal is proportional to substrate phosphorylation. Assays were performed in white 96-well low-volume HTRF microplates (Revvity). Recombinant full-length human TBK1 was purchased from Promega (#V3991). Reactions were carried out in KinEASE kinase buffer supplemented with 5 mM MgCl_2_ and 1 mM DTT. Ap_4_A (Jena Bioscience) was serially diluted 1:2 in kinase buffer containing 1.25 µM STK substrate-biotin and 125 µM ATP. Eleven Ap_4_A-containing dilution steps were prepared starting from 8 mM Ap_4_A; a twelfth condition lacking Ap_4_A served as the uninhibited control. Eight microliters of each Ap_4_A/substrate/ATP mixture were transferred to the assay plate, and reactions were initiated by adding 2 µL TBK1 diluted in the same kinase buffer. Final reactions contained 1 ng/µL TBK1, 1 µM STK substrate-biotin, 100 µM ATP, and Ap_4_A concentrations ranging from 6.4 mM to 0.

Reactions were incubated for 1 h at room temperature in sealed plates and stopped by addition of 10 µL detection reagent containing STK antibody-cryptate, streptavidin-XL665, and EDTA in KinEASE detection buffer. Streptavidin-XL665 was used at a final assay concentration of 62.5 nM, corresponding to the recommended 8:1 biotin/streptavidin ratio for 1 µM substrate in the enzymatic reaction. Plates were sealed again and incubated for 1 h at room temperature before readout.

HTRF signals were measured on a Spark 20M multimode plate reader equipped with an HTRF module (Tecan). Sequential donor and acceptor emission measurements were performed at 620 nm and 665 nm, respectively, using excitation at 320 nm/25 nm, acceptor emission at 665 nm/8 nm, donor emission at 620 nm/10 nm, 50 flashes, a 100 µs lag time, a 300 µs integration time, a 510 nm dichroic mirror, and optimized gain settings.

The HTRF ratio was calculated as the 665 nm/620 nm signal multiplied by 10,000. Background-corrected signals were obtained by subtracting the ratio of negative-control reactions lacking kinase from the corresponding kinase-containing reactions. Each experiment was performed independently three times. The dose-response curve was fitted in GraphPad Prism using nonlinear regression with a four-parameter variable-slope inhibitor-response model ([inhibitor] vs. response). Fit was obtained by least-squares regression. The goodness of fit is reported as R^2^, and inhibitory potency is reported as the D/S_50_, defined as the molar Ap_4_A:ATP ratio required to reduce TBK1 activity by 50 %.

### Preparation of CRISPR/Cas9 plasmids

NUDT2 knockout and scrambled sgRNA control THP-1 cells were generated using the lentiviral CRISPR/Cas9 vector pL-CRISPR.EFS.PAC (Addgene plasmid #57828, RRID:Addgene_57828) (Heckl et al., 2014). This vector encodes *Sp*Cas9, an sgRNA expression cassette, and a puromycin-resistance marker. The NUDT2-targeting sgRNA was designed using the Synthego design tool, prioritizing predicted on-target activity, low predicted off-target activity, suitable GC content, and proximity to the NUDT2 start codon. The NUDT2-targeting sgRNA sequence was 5′-GGA GAG GAT GAC TTG GAA AC-3′. The scrambled control sgRNA sequence was 5′-GCA CTC ACA TCG CTA CAT CA-3′.Oligonucleotides encoding the sgRNA were synthesized by Sigma-Aldrich with *BsmBI*-compatible overhangs and subsequently cloned into pL-CRISPR.EFS.PAC. Final plasmids were verified by Sanger sequencing performed by Microsynth.

### Production of lentiviral particles

Lentiviral particles were produced in HEK293T cells using the PEI-based Transporter 5 Transfection Reagent (Kyfora Bio). The day before transfection, 5 x 10^6^ HEK293T cells were cultured in a 10-cm cell culture dish in 12 mL complete DMEM containing 10 % FCS, 1 % L-glutamine, 1 % penicillin/streptomycin, 1 % sodium pyruvate, and 1 % MEM non-essential amino acids, at 37 °C and 5 % CO_2_.

On the day of transfection, the medium was replaced with 6 mL DMEM containing 2 % FCS and 0.1 % penicillin/streptomycin, while all other supplements were kept unchanged. For each viral production, 6 µg plasmid, 10 µg psPAX2 packaging plasmid (Addgene #12260; RRID:Addgene_12260), and 2 µg pMD2.G envelope plasmid (Addgene #12259; RRID:Addgene_12259) were diluted in 150 mM NaCl to a final volume of 600 µL. The mixture was vortexed briefly and incubated for 5 min at room temperature. Subsequently, 59.4 µL Transporter 5 Transfection Reagent was added, mixed by gentle inverting, and incubated for 30 min at room temperature to allow DNA-PEI complex formation.

The transfection mixture was added dropwise to the HEK293T cells and distributed by gentle swirling. Cells were incubated for 6 h at 37 °C and 5 % CO_2_. The medium was then removed and replaced with 10 mL complete DMEM supplemented with 2 mM caffeine. Viral supernatant was harvested 48 and 72 h after transfection, sterile-filtered through a 0.45 µm syringe filter (Sarstedt), and stored at −80 °C until use. HEK293T cells (DSMZ ACC-635; RRID:CVCL_0063) were obtained from Leibniz-Institute, Deutsche Sammlung von Mikroorganismen und Zellkulturen (DSMZ) and maintained as low-passage cultures. Cells were regularly tested for mycoplasma contamination.

### Lentiviral transduction and generation of THP-1 single-cell clones

THP-1 cells were transduced with lentiviral particles carrying either the NUDT2-targeting sgRNA or the scrambled sgRNA control construct. For transduction, THP-1 cells were seeded in 6-well plates coated with RetroNectin (Takara Bio) at 2 x 10^5^ cells per well in 2 mL complete RPMI 1640 medium containing 10 % FCS and 1 % penicillin/streptomycin. Viral supernatant harvested 48 and 72 h after HEK293T transfection wer thawed on ice, concentrated using Lenti-X concentrator (Takara Bio) and added to the cells at 1 mL per well together with polybrene at a final concentration of 8 µg/mL. Cells were centrifuged at 2,000 rpm for 90 min at 32 °C and then incubated overnight at 37 °C and 5 % CO_2_. The following day, cells were pelleted by centrifugation at 300 x *g* for 5 min, resuspended in fresh complete RPMI medium, and returned to culture.

Three days after transduction, puromycin selection was started by culturing cells in complete RPMI medium containing 1.5 µg/mL puromycin. Selection was continued until non-transduced control cells were eliminated and surviving transduced cells had recovered. Puromycin-resistant NUDT2-targeted and scrambled gRNA control bulk populations were then used for single-cell isolation.

For clonal isolation, viable cells were selected by FACS using forward- and side-scatter gating and seeded as single cells into U-bottom 96-well plates. Parental THP-1 cells were processed in parallel by the same strategy.

Single-cell status, colony formation, and culture health were monitored microscopically throughout clonal outgrowth. Clones were expanded stepwise from U-bottom 96-well plates to larger culture formats. Transfer to larger culture vessels was performed according to cell density, viability, and overall culture health.

Established parental, scrambled gRNA control, and NUDT2-knockout single-cell clones were cryopreserved in 90 % FCS and 10 % DMSO and slowly frozen to −80 °C. THP-1 cells (DSMZ ACC-16; RRID:CVCL_0006) were obtained from DSMZ and maintained as low-passage cultures. Cells were regularly tested for mycoplasma contamination.

### Extraction of intracellular nucleotides from THP-1 single-cell clones

Individual THP-1 single-cell clones were thawed from cryopreserved stocks into Gibco™ RPMI 1640 medium supplemented with 10 % (v/v) FCS and 1 % (v/v) penicillin/streptomycin. Cells were maintained in culture for 6−8 days to allow full recovery, until they reached ≥95 % viability and displayed a healthy suspension-culture phenotype without visible debris or clumping. Cells were then expanded for 48 h before nucleotide extraction.

Cell density, viability, and diameter were determined using a LUNA-FL™ dual fluorescence cell counter (Logos Biosystems) with acridine orange/propidium iodide staining (Logos Biosystems, F23001). Briefly, 18 µL cell suspension was mixed with 2 µL staining solution, loaded onto LUNA™ cell counting slides (Logos Biosystems, L12001), and analyzed according to the manufacturer’s instructions.

For nucleotide analysis, 1 x 10^6^ cells were collected per extraction replicate in 2 mL microcentrifuge tubes. Two independent extraction aliquots were prepared from each single-cell clone. Cell aliquots were pelleted by centrifugation at 300 x *g* for 5 min using a swing-out rotor, and the supernatant was carefully removed. Pellets were washed once with 900 µL PBS (Carl Roth), centrifuged again, and the supernatant was removed as completely as possible without disturbing the pellet.

Nucleotides were extracted by adding 100 µL extraction buffer (pre-cooled to −20 °C) containing 5 mM Tris, 0.5 mM EDTA, and 50 % (v/v) methanol, pH 7.2, followed by 100 µL chloroform (pre-cooled to −20 °C). Samples were vigorously shaken for 30 min at −20 °C and then centrifuged at 10,000 x *g* for 10 min at −9 °C using a fixed-angle rotor to separate the phases. The upper polar phase was carefully collected and filtered through a 0.22 µm PTFE membrane filter with 4 mm diameter (Phenomenex) into fresh LC–MS sample vials (Macherey-Nagel).

For filtration, a 1 mL single-use syringe was prepared by removing the plunger and attaching the PTFE membrane filter. The collected upper polar phase was transferred into the syringe using a pipette, the plunger was reinserted, and the extract was pushed through the filter. Extracted samples were stored at −80 °C until LC-QQQ-MS analysis.

### Quantitative determination of target analytes was performed using a LC-MS/MS

The chromatographic separation was performed on an Agilent Infinity II 1290 HPLC system using a SeQuant ZIC-pHILIC column (150 x 2.1 mm, 5 μm particle size, peek coated, Merck) connected to a guard column of similar specificity (20 x 2.1 mm, 5 μm particle size, Phenomoenex) a constant flow rate of 0.1 ml/min with mobile phase A comprised of 10 mM ammonium acetate in water, pH 9, supplemented with medronic acid to a final concentration of 5 μM and mobile phase B comprised of 10 mM ammonium acetate in 90:10 acetonitrile to water, pH 9 at 40° C. The injection volume was 2 µl. The mobile phase profile consisted of the following steps and linear gradients: 0−1 min constant at 75 % B; 1−6 min from 75 to 40 % B; 6 to 9 min constant at 40 % B; 9−9.1 min from 40 to 75 % B; 9.1 to 20 min constant at 75 % B. An Agilent 6495 ion funnel mass spectrometer was used in negative mode with an electrospray ionization source and the following conditions: ESI spray voltage 3500 V, nozzle voltage 1000 V, sheath gas 300° C at 9 l/min, nebulizer pressure 20 psig and drying gas 100° C at 11 l/min. Compounds were identified based on their mass transition and retention time compared to standards. Chromatograms were integrated using MassHunter software (Agilent, Santa Clara, CA, USA

Absolute concentrations were determined based on an external standard curve covering a concentration range of 0.1−1000 µM for AMP, ADP, ATP, GMP, GDP, and GTP (Jena Bioscience) and of 0.01−100 µM for Ap_3_A, Ap_4_A and Ap_4_G (Jena Bioscience). Calibration curves were fitted with a power-function regression model, and all accepted curves had R^2^ values ≥ 0.998. Mass transitions, collision energies, cell accelerator voltages, and dwell times have been optimized using chemically pure standards. The parameter settings of all targets are given in table S3.1.

To estimate apparent intracellular nucleotide concentrations, the average cell diameter was monitored and measured using a LUNA-FL™ cell counter. This measurement accounts for variation in cell diameter within each cell population by calculating the mean diameter of the analyzed cells. Cell diameter did not substantially differ between clones; therefore, the average diameter across all measurements was used for cell-volume calculations. This average diameter was 14.5 µm. Assuming spherical cells, the mean single-cell volume was calculated from this diameter. Intracellular nucleotide concentrations were then calculated from the measured nucleotide amounts in the 100 µL extraction volume and normalized to the total calculated cell volume of the extracted cells.

### Immunoblotting and signaling analysis

To validate the NUDT2 status of THP-1 single cell clones we used western blotting to detect NUDT2. Whole-cell lysates were prepared from THP-1 parental clones (n = 6), sgRNA scramble clones (n = 6), and Nudt2-knockout clones (n = 6). Cells were lysed for 60 min on ice with lysis buffer (50 mM Tris-HCl, pH 7.5, 150 mM NaCl, 1 % Triton X-100, 1:10 protease inhibitor cocktail (Sigma-Aldrich)) and clarified by centrifugation at 13,000 rpm for 10 min. Equal amounts of protein (5 µg) were separated by 10% SDS-PAGE and transferred onto methanol-activated PVDF membranes using a semi-dry transfer system (BioRad). Membranes were blocked in TBST containing 10 % milk for 1 h at room temperature and incubated overnight at 4 °C with primary antibodies against human NUDT2 (Abcam, #ab210769, 1:5,000) or β-Actin (Sigma-Aldrich, #A1978, RRID:AB_476692, 1:10,000). After washing, membranes were incubated with HRP-conjugated secondary antibodies against rabbit IgG (Cell Signaling Technology, #7074, RRID:AB_2099233, 1:20,000) or mouse IgG (Agilent Dako, #P044701-2 / P0447, RRID:AB_2617137, 1:20,000), respectively. Bound antibodies were detected using ECL Prime Western Blotting Detection Reagent (Cytiva, #RPN2236; solutions 1 and 2 mixed 1:1) and imaged using a c400 Imaging System (Azure Biosystems).

## Data availability

Protein structure coordinates and structure factors have been deposited in the Protein Data Bank in Europe (PDBe) under accession codes 30LM (ATP-bound *Mm* cGAS) and 30LN (pppGpA-bound *Mm* cGAS). Source data underlying the quantitative plots are provided with the manuscript. All other data generated or analyzed in this study are available upon reasonable request.

## Disclosure

The authors declare no conflict of interests. Generative language AI tools, including ChatGPT- 5.5 (OpenAI) and Perplexity, were used during manuscript preparation to assist with language editing, grammar, readability, and textual consistency. The authors reviewed all AI-assisted revisions and take full responsibility for the scientific content of the manuscript.

## Article and author information

### Author details

**Paul Weiland**

Department of Chemistry & Center for Synthetic Microbiology (SYNMIKRO), Marburg University, Marburg, Germany and Department of Medicine & Center for Tumor Biology and Immunology (ZTI), Marburg University, Marburg, Germany

**Contribution**

Conceptualization, Data curation, Formal analysis, Investigation, Methodology, Project administration, Validation, Visualization, Writing – original draft, Writing – review and editing

**Competing interests**

No competing interests declared

**Renuka Dharani Shivakumar**

Department of Chemistry & Center for Synthetic Microbiology (SYNMIKRO), Marburg University, Marburg, Germany

**Contribution**

Formal analysis, Investigation, Visualization, Writing – review and editing

**Competing interests**

No competing interests declared

**Julia Schmidt**

Department of Chemistry & Center for Synthetic Microbiology (SYNMIKRO), Marburg University, Marburg, Germany

**Contribution**

Investigation, Visualization

**Competing interests**

No competing interests declared

**Ekaterina Jalomo-Khayrova**

Department of Chemistry & Center for Synthetic Microbiology (SYNMIKRO), Marburg University, Marburg, Germany

**Contribution**

Formal analysis, Investigation, Visualization

**Competing interests**

No competing interests declared

**Victor Zegarra**

Department of Chemistry & Center for Synthetic Microbiology (SYNMIKRO), Marburg University, Marburg, Germany

Max Planck Institute for Biological Intelligence, Munich, Germany

**Contribution**

Investigation, Formal analysis

**Competing interests**

No competing interests declared

**Ying Wang**

Department of Medicine & Center for Tumor Biology and Immunology (ZTI), Marburg University, Marburg, Germany

**Contribution**

Investigation

**Competing interests**

No competing interests declared

**Nicole Paczia**

Max Planck Institute for Terrestrial Microbiology, Marburg, Germany

**Contribution**

Formal analysis, Methodology, Resources, Writing – review and editing

**Competing interests**

No competing interests declared

**Stephan Kiontke**

Department of Chemistry & Center for Synthetic Microbiology (SYNMIKRO), Marburg University, Marburg, Germany

**Contribution**

Data curation, Formal analysis, Writing – review and editing

**Competing interests**

No competing interests declared

**Andreas Burchert**

Department of Medicine & Center for Tumor Biology and Immunology (ZTI), Marburg University, Marburg, Germany; Department of Hematology, Oncology and Immunology, University Hospital Giessen and Marburg, Marburg, Germany

**Contribution**

Funding acquisition, Project administration, Resources, Supervision, Writing – review and editing

**Competing interests**

No competing interests declared

**Gert Bange**

Department of Chemistry & Center for Synthetic Microbiology (SYNMIKRO), Marburg University, Marburg, Germany; Max Planck Institute for Terrestrial Microbiology, Marburg, Germany

**Contribution**

Conceptualization, Funding acquisition, Project administration, Resources, Supervision, Validation, Writing – review and editing

**Competing interests**

No competing interests declared

## Acknowledgements

We thank the beamline staff at ESRF (Grenoble, France) and DESY (Hamburg, Germany) for their support during X-ray diffraction data collection. We thank Peter Claus from the Metabolomics Facility at the Max Planck Institute for Terrestrial Microbiology, Marburg, for assistance with QQQ-MS measurements. Additionally, we gratefully acknowledge the support and resources provided by the ChemBio research-based platform for assistance with establishing the HTRF assays and Prof. Dr. Olalla Vázquez for access to the HTRF-compatible plate reader.

---

**Figure S1.1.**
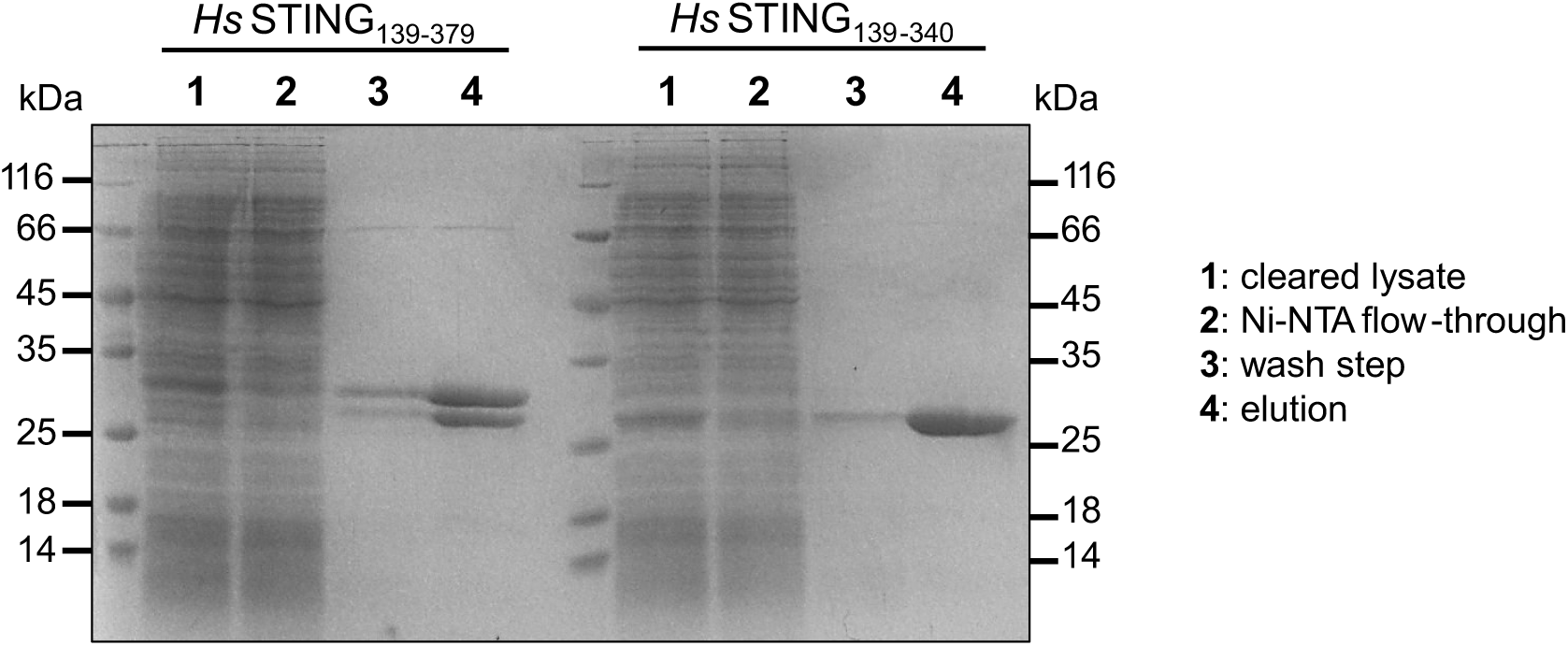
SDS-PAGE analysis of recombinant human STING constructs showing C-terminal heterogeneity. SDS-PAGE analysis of purified full-length cytosolic human STING_139−379_ following recombinant expression and Ni-NTA affinity purification. In addition to the expected full-length species, a lower molecular weight band is observed, consistent with partial C-terminal truncation corresponding to a ΔC-terminal STING_139−340_ species (Fig. 1 A). This heterogeneity prompted the use of human STING constructs lacking the C-terminal tail for subsequent binding studies.

**Figure S1.2.**
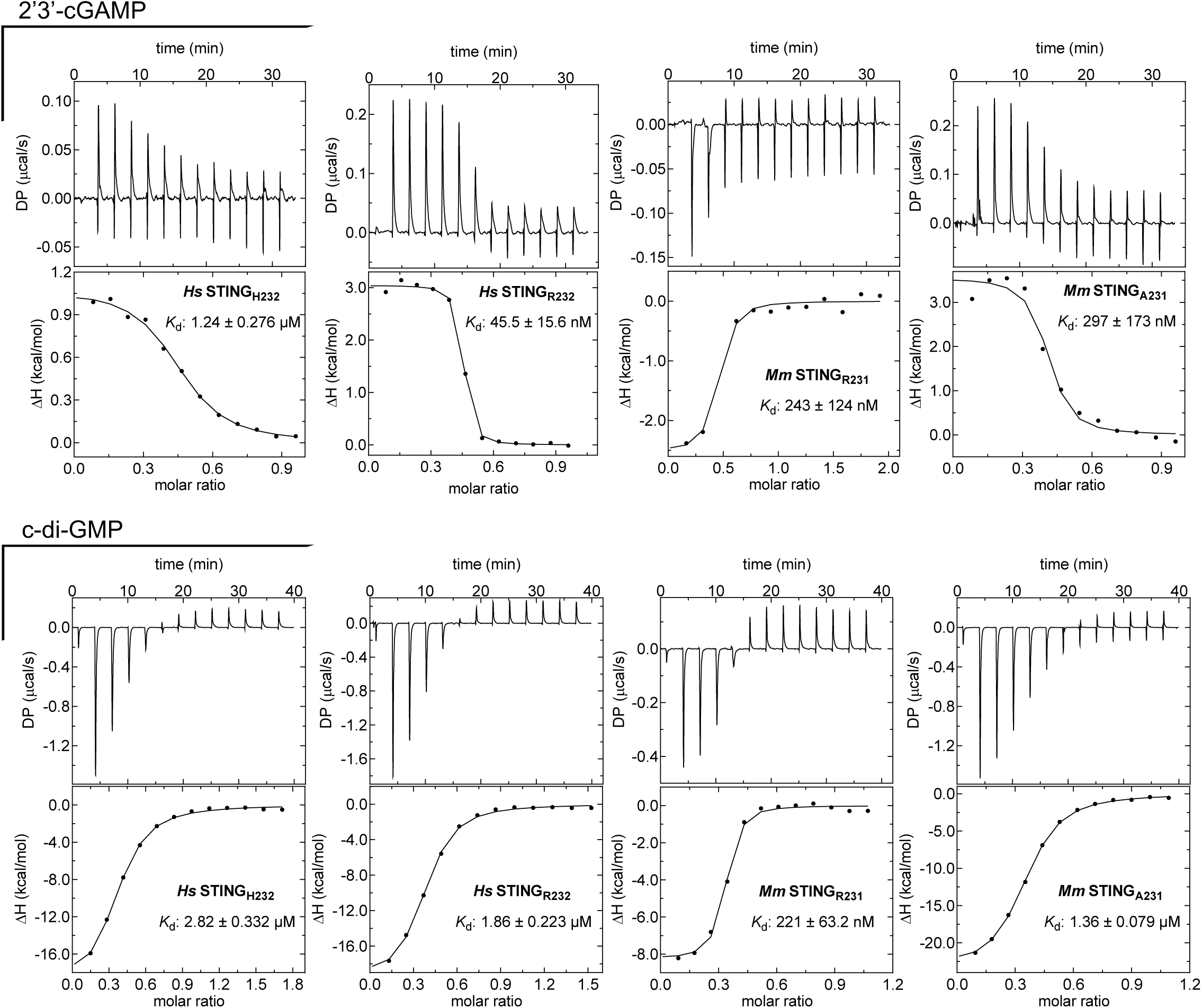
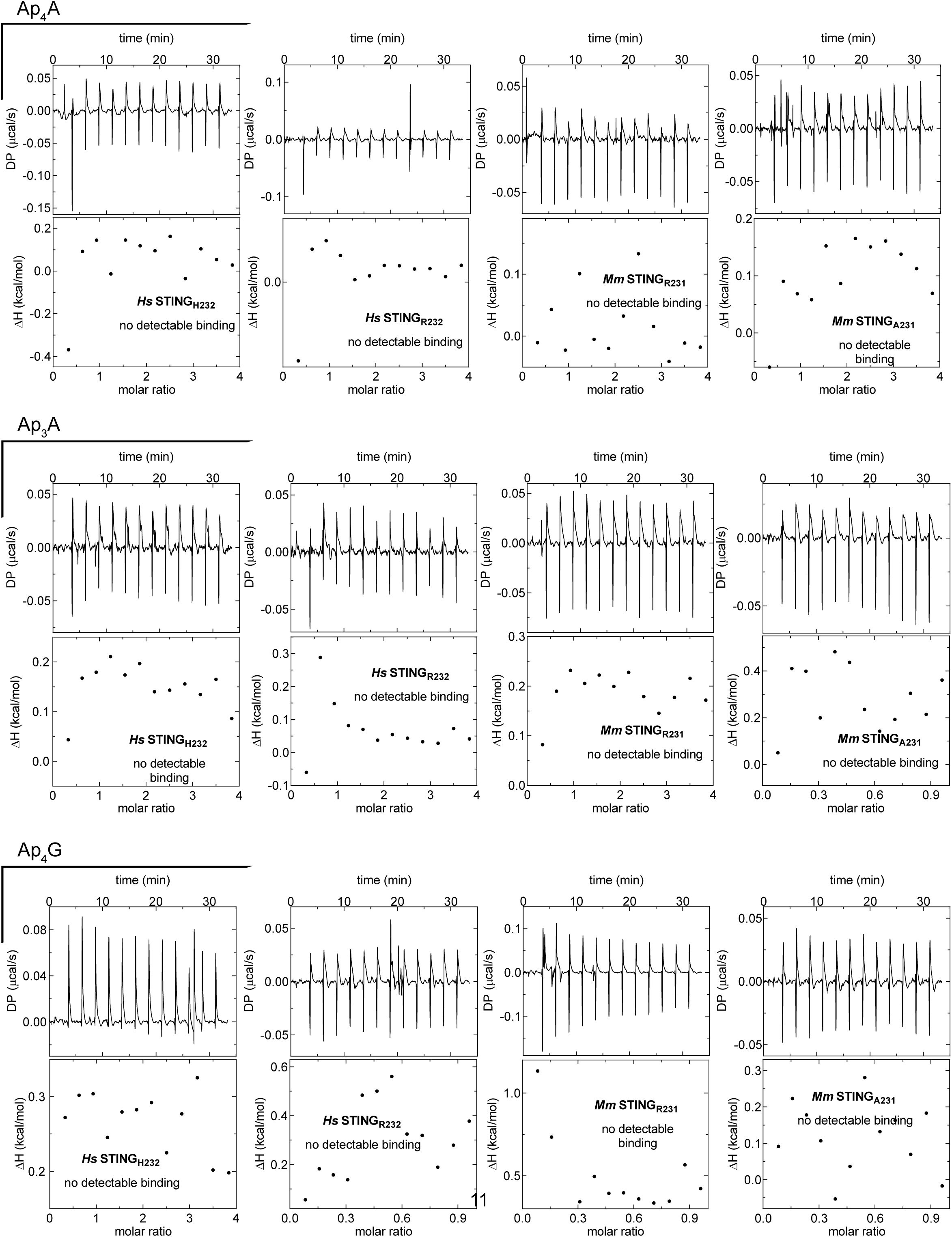
Representative ITC datasets for STING binding. Shown are representative ITC replicates for binding experiments between STING constructs and canonical ligands (2′3′-cGAMP and c-di-GMP as positive controls) or dinucleoside polyphosphates (Ap_4_A, Ap_3_A, and Ap_4_G). Raw thermograms (top) and integrated binding isotherms (bottom) are shown, demonstrating robust binding of canonical ligands and absence of detectable binding for dinucleoside polyphosphates.

**Figure S1.3.**
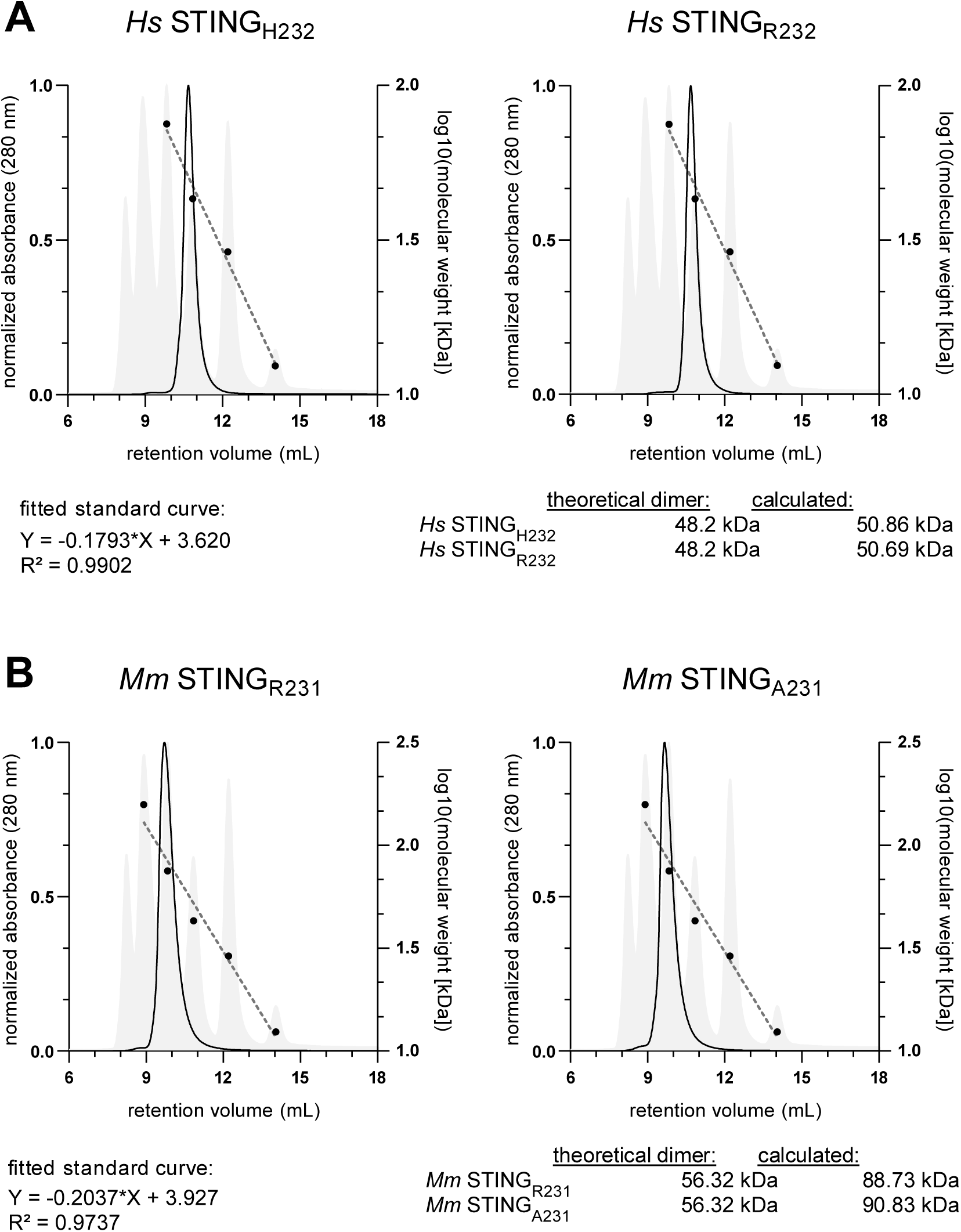
Analytical SEC validation of STING oligomeric state. Analytical Size-Exclusion Chromatography (SEC) of human and mouse STING constructs used for binding experiments. The column was calibrated using globular protein standards of known molecular weight: cytochrome c (12.4 kDa, Sigma-Aldrich), carbonic anhydrase (29 kDa, Cytiva), ovalbumin (43 kDa, Cytiva), conalbumin (75 kDa, Cytiva) and aldolase (158 kDa, Cytiva). Calculated molecular weights of recombinant STING constructs correspond to the expected theoretical dimeric assemblies, confirming that all constructs are predominantly dimeric in solution under the conditions used. The mouse constructs eluted at a slightly higher apparent molecular weight, likely reflecting altered migration caused by their retained, disordered C-terminal tails.

**Figure S1.4.**
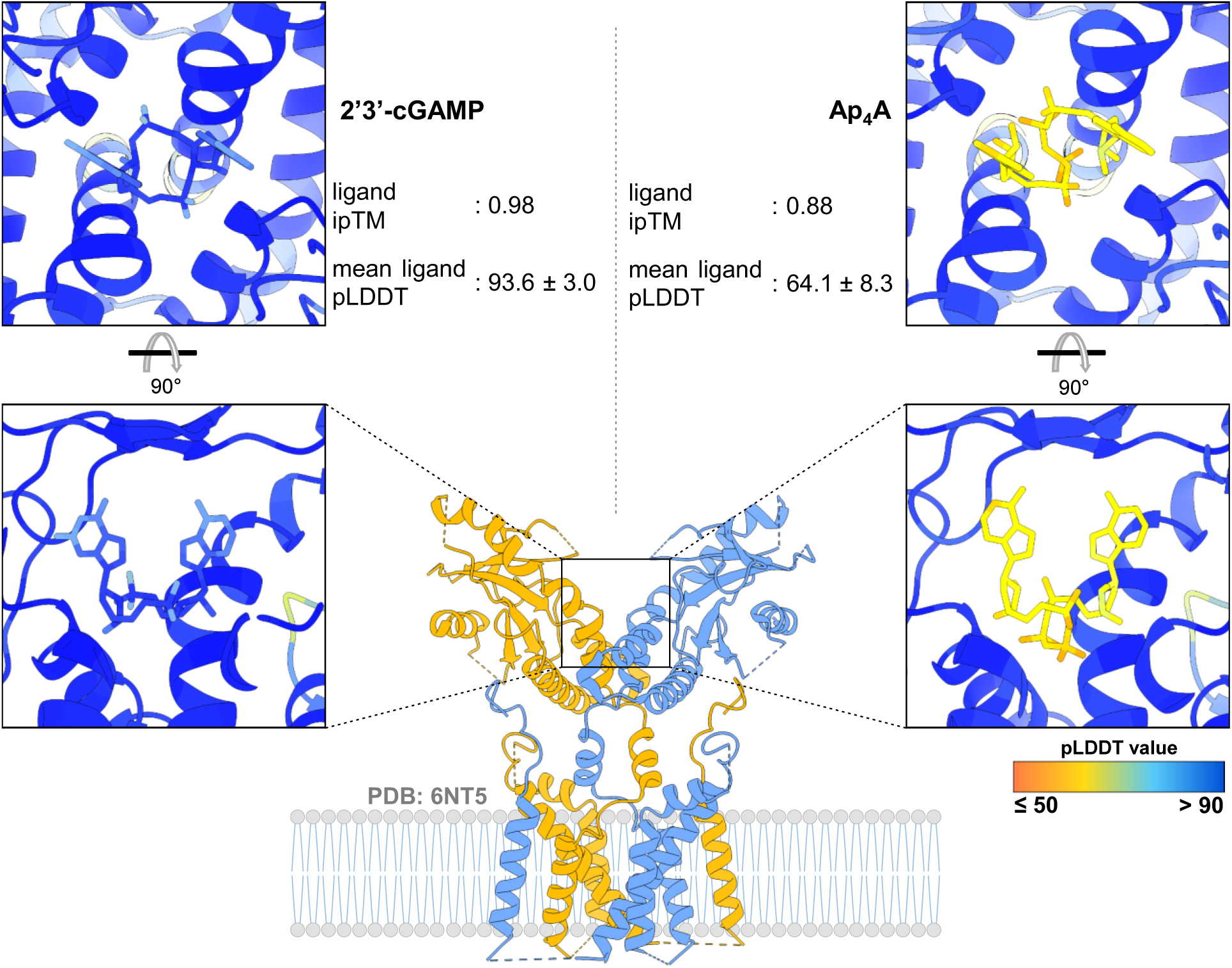
Predicted structural constraints on Ap_4_A binding within the STING dimer. Structural representations of the STING dimer (PDB: 6NT5) highlighting the ligand-binding pocket in the context of the dimeric assembly (box, image center). Boltz2 models illustrate spatial constraints imposed by dimerization that limit accommodation of larger dinucleotides such as Ap_4_A (right side), in contrast to canonical cyclic dinucleotide ligands like 2′3′-cGAMP (left side).

**Figure S2.1.**
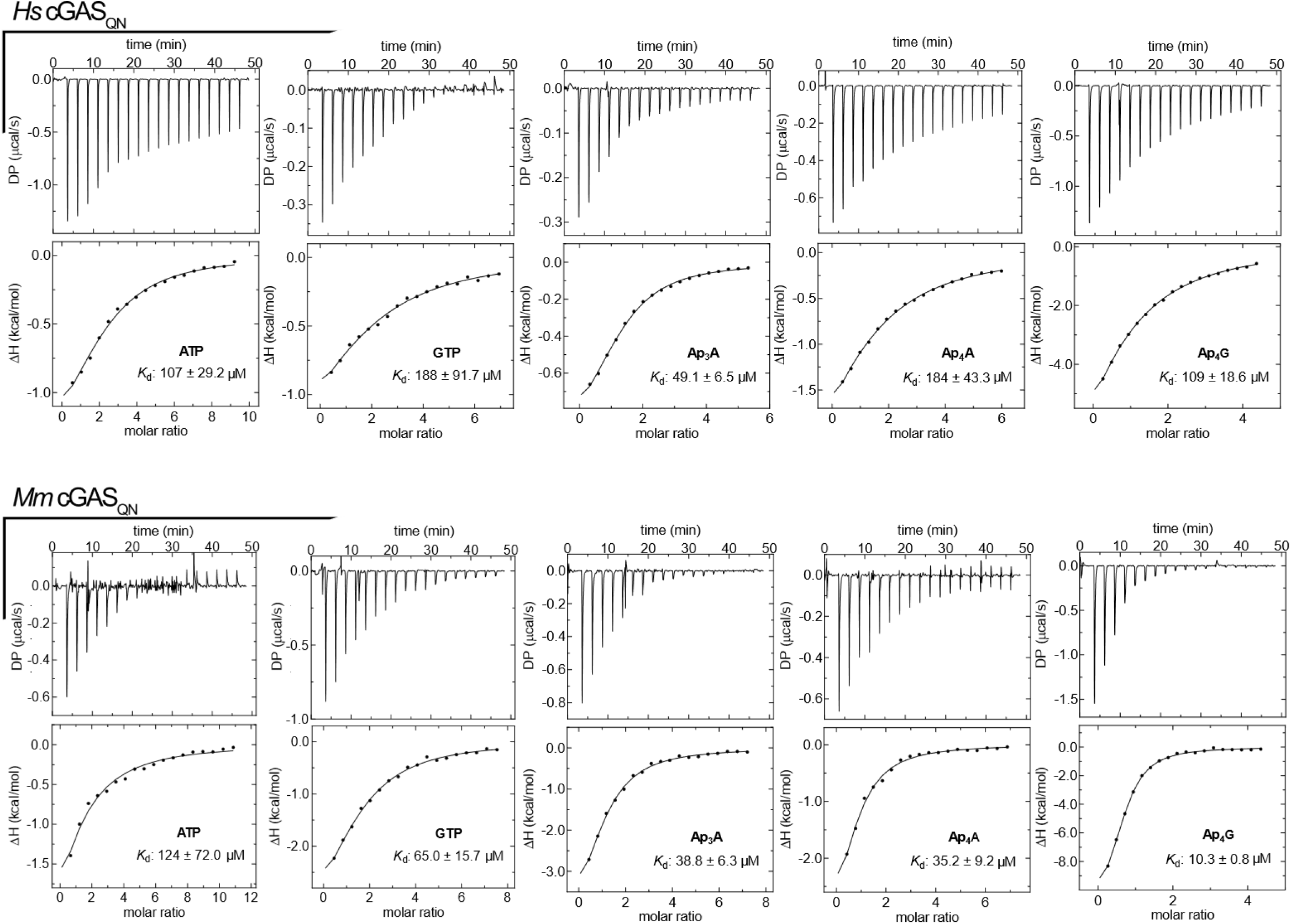
Representative ITC datasets for cGAS binding in absence of DNA. Shown are representative ITC replicates for nucleotide binding to human and mouse cGAS_QN_ in the absence of DNA. Raw thermograms (top) and integrated binding isotherms (bottom) for ATP, GTP, Ap_3_A, Ap_4_A, and Ap_4_G. All measurements were performed under identical buffer conditions.

**Figure S2.2.**
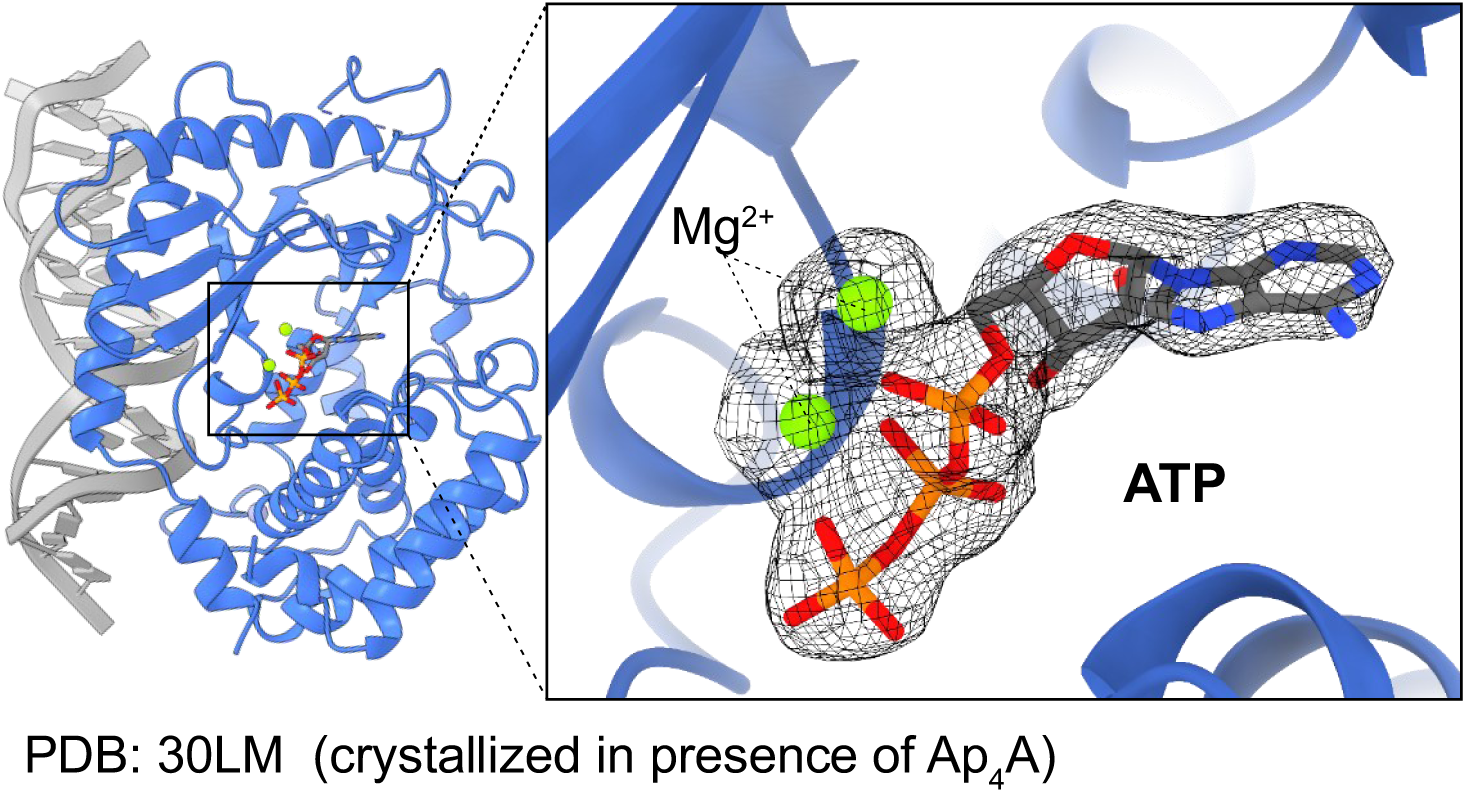
Electron density in nucleotide binding site of DNA-bound cGAS crystallized with Ap_4_A. Electron density observed in the active site of DNA-bound mouse cGAS_147−507_ crystallized in the presence of Ap_4_A. Because the observed electron density was only consistent with ATP and not with intact Ap_4_A, ATP was modeled in the binding pocket and is shown with the corresponding 2*F*_o_-*F*_c_ map contoured at 1 σ shown in black. The structure was deposited in the Protein Data Bank under accession code 30LM.

**Figure S2.3.**
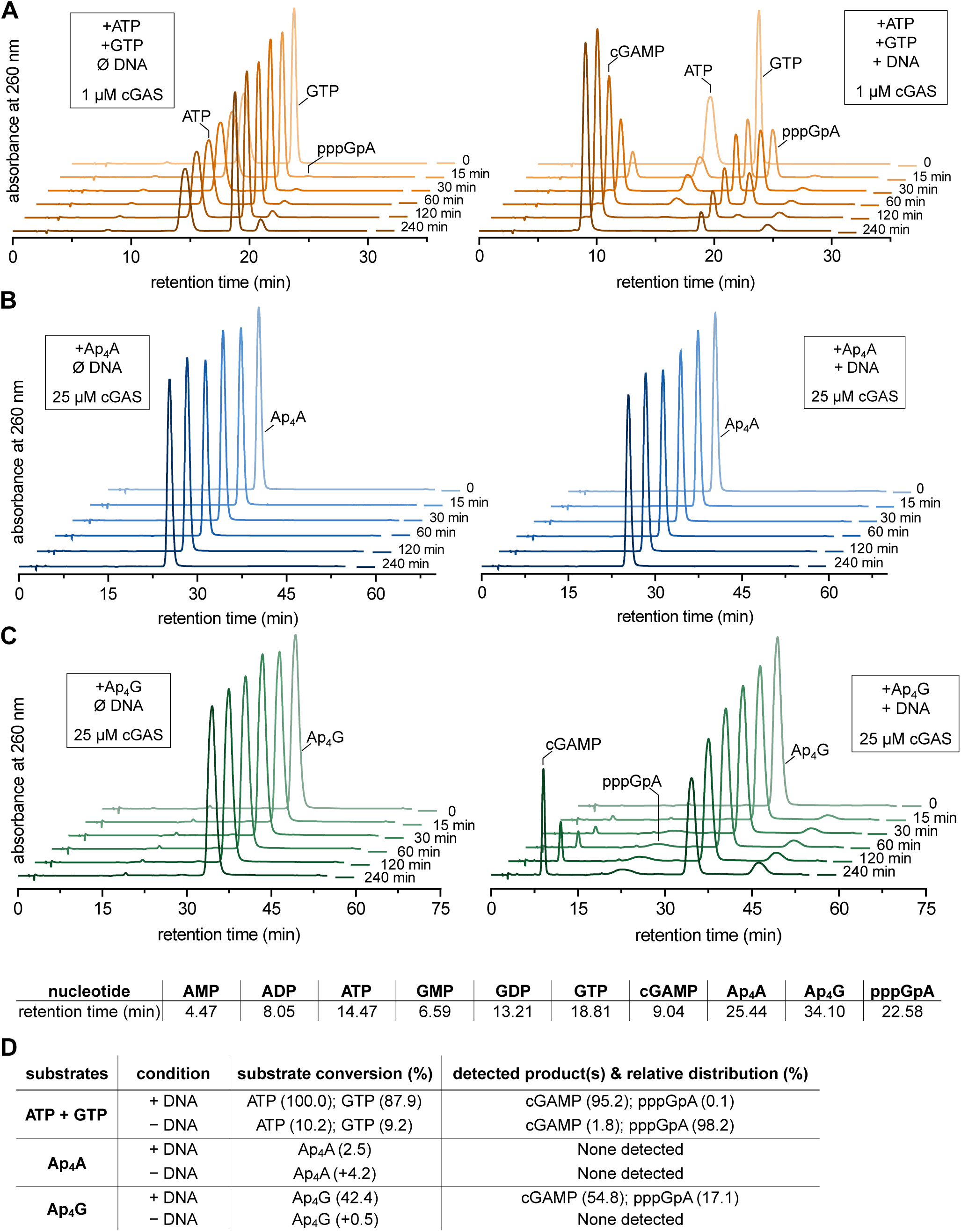
Full HPLC time courses of cGAS reaction products. [**A**−**C**] HPLC analysis of cGAS reaction products generated from canonical ATP/GTP substrates or dinucleoside polyphosphates in the absence or presence of 60-bp dsDNA. Canonical reactions were performed with 1 µM mouse cGAS_147−507_, whereas Ap_4_A and Ap_4_G reactions were performed with 25 µM to maximize detection of low-efficiency turnover. Reactions were sampled after 0, 15, 30, 60, 120, and 240 min. Ap_4_A remained largely unchanged and did not support detectable cGAMP formation either in the absence or presence of DNA. Ap_4_G remained largely unchanged in the absence of DNA but yielded detectable cGAMP and pppGpA in the presence of DNA, indicating inefficient DNA-dependent processing of Ap_4_G. Retention times of nucleotide standards used for peak assignment are shown in the table below. Chromatograms were recorded by absorbance at 260 nm. [**D**]

**Figure S3.1.**
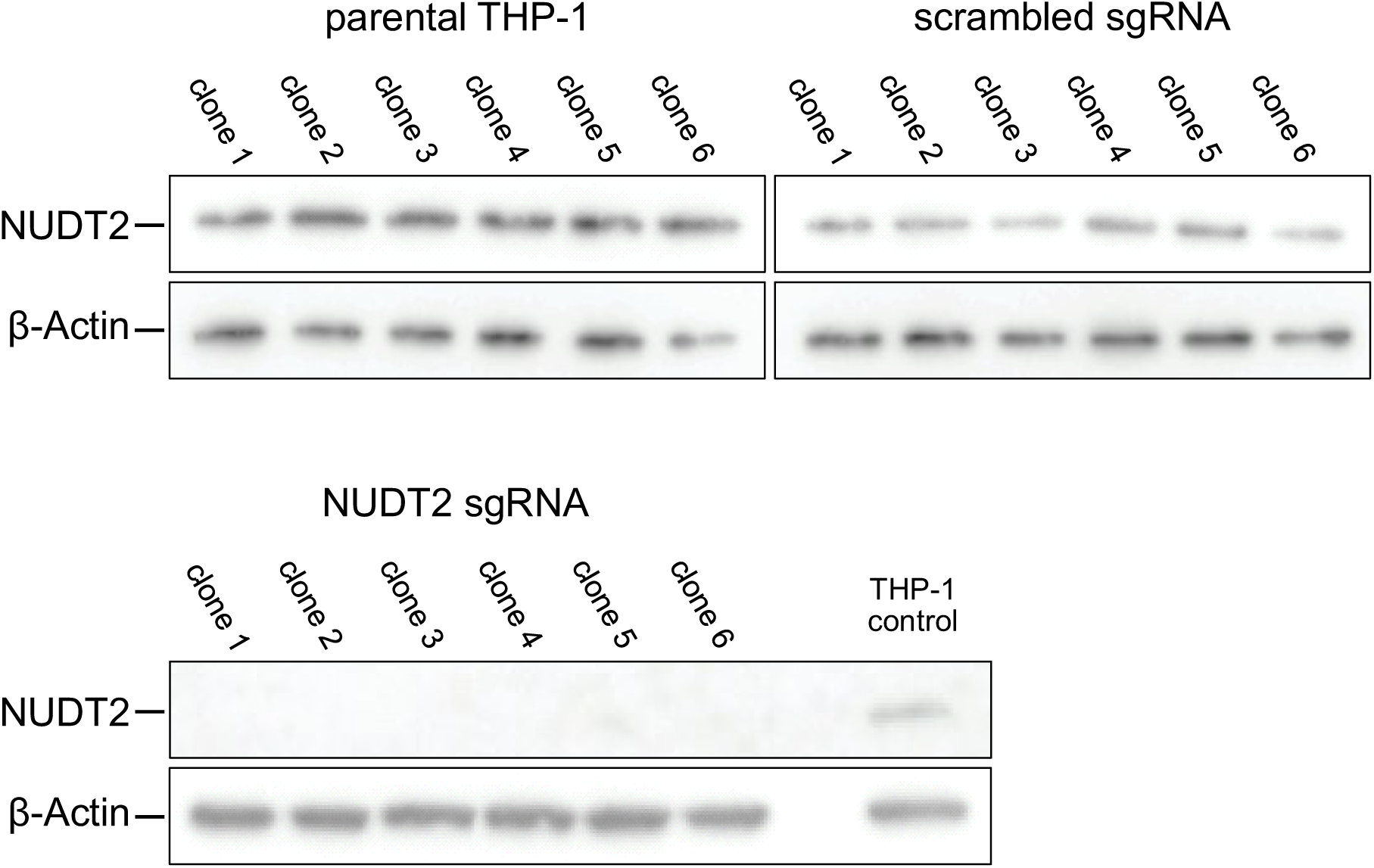
Western blot validation of NUDT2 status in THP-1 cells. Western blot analysis of NUDT2 protein levels in THP-1 parental clones, scrambled sgRNA control clones, and NUDT2 sgRNA knockout clones. Loss of NUDT2 protein confirms successful genetic disruption in the knockout cell clones. A representative loading control is shown (β-Actin).

**Figure S3.2.**
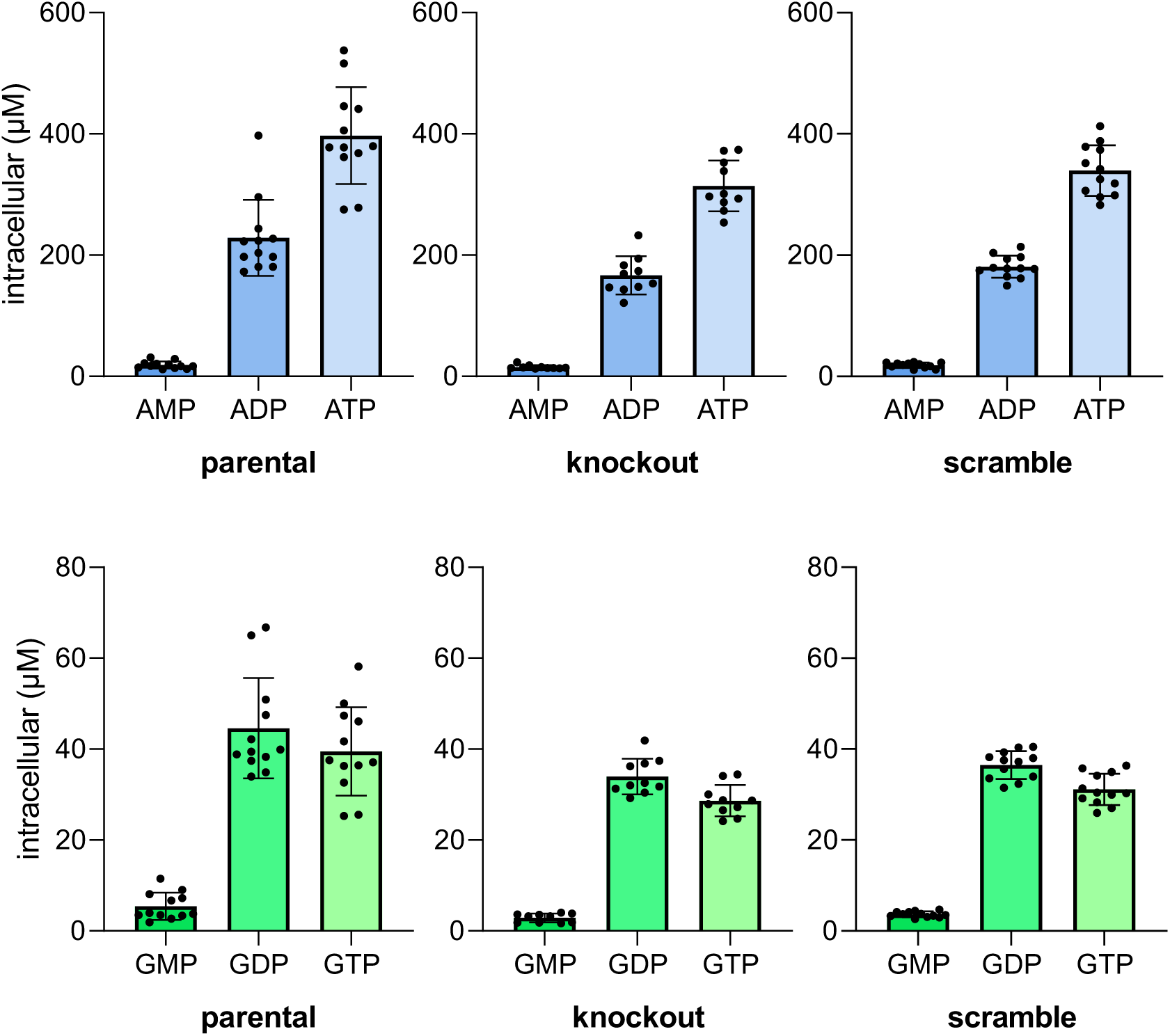
**Intracellular adenine and guanine nucleotide pools in THP-1 cells**LC-QQQ-MS analysis of intracellular adenine and guanine nucleotides in parental, scrambled gRNA control, and NUDT2 knockout THP-1 cells. Shown are the intracellular concentrations of AMP, ADP, ATP, GMP, GDP, and GTP. Across all three cell lines, canonical adenine and guanine nucleotide pools were not significantly altered by NUDT2 loss, providing a stable reference for comparison with dinucleoside polyphosphate accumulation. Parental and scrambled sgRNA control cells were represented by six independent single-cell clones and NUDT2-knockout cells by five; each clone was extracted in duplicate. Bars represent means with standard deviations and individual points represent duplicate extracts from independent single-cell clones.

**Table S1.1.**
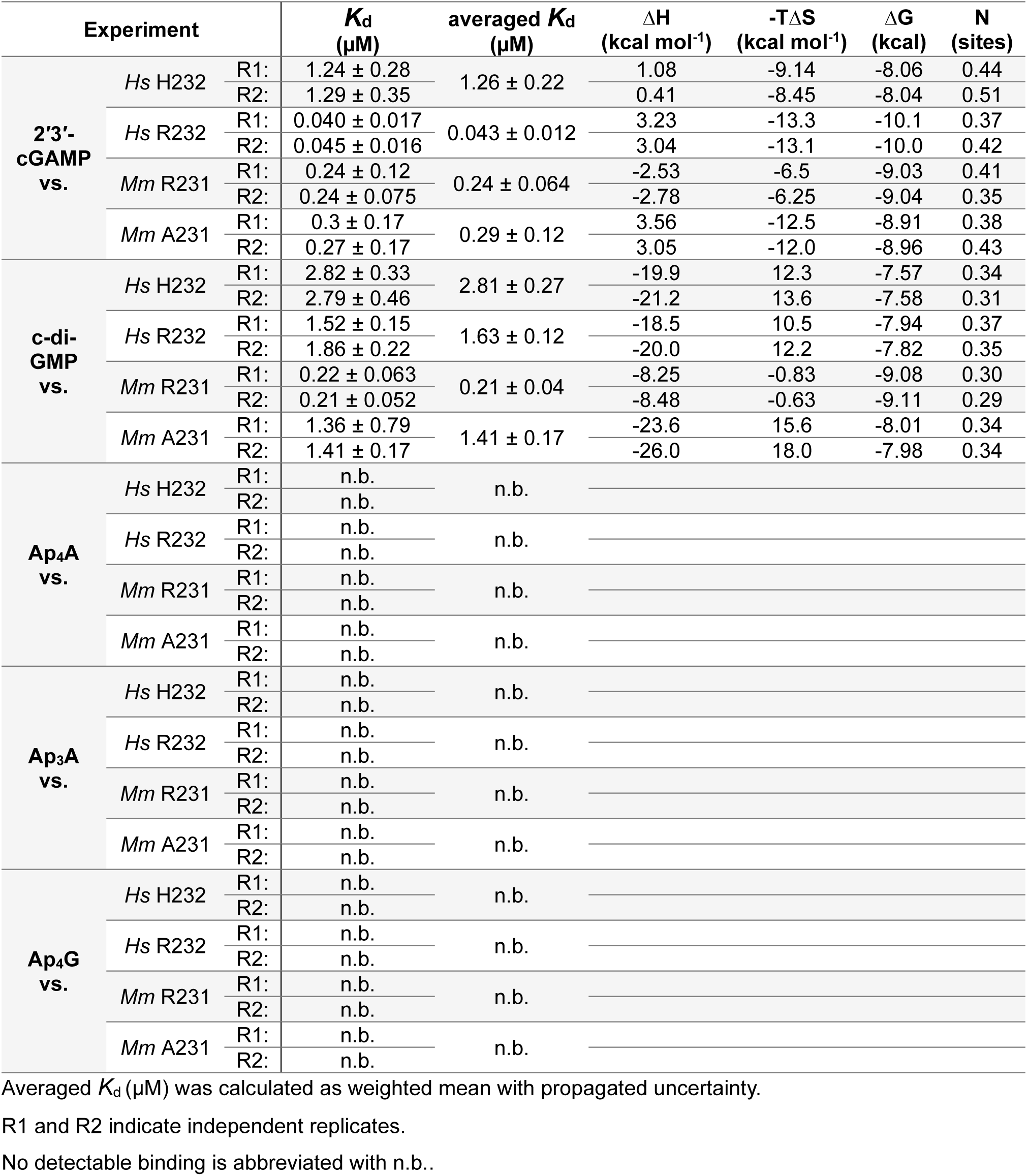
Summarized thermodynamic values determined from ITC for STING. Thermodynamic values determined from all ITC experiments of STING constructs against different nucleotides.

**Table S2.1.** Summarized thermodynamic values determined from ITC for cGAS. Thermodynamic values determined from all ITC experiments of cGAS constructs against different nucleotides in absence of DNA.

| Experiment | | | $K_d$<br>($\mu\text{M}$) | averaged $K_d$<br>($\mu\text{M}$) | $\Delta H$<br>(kcal mol<sup>-1</sup>) | $-T\Delta S$<br>(kcal mol<sup>-1</sup>) | $\Delta G$<br>(kcal) | N<br>(sites) |
| --- | --- | --- | --- | --- | --- | --- | --- | --- |
| **GTP**<br>**vs.** | *Hs* cGAS | R1: | 188 ± 91.7 | 167.6 ± 42.4 | -1.96 ± 0.8 | -3.13 | -5.09 | 1.90 |
|  |  | R2: | 162 ± 47.8 |  | -1.84 ± 0.6 | -3.56 | -5.21 | 1.82 |
|  | *Mm* cGAS | R1: | 68.8 ± 13.3 | 67.2 ± 10.1 | -8.61 | 2.93 | -5.68 | 1.03 |
|  |  | R2: | 65.0 ± 15.7 |  | -4.51 | -1.20 | -5.71 | 1.52 |
| **ATP**<br>**vs.** | *Hs* cGAS | R1: | 70.8 ± 24.3 | 85.6 ± 18.7 | -0.86 | -4.80 | -5.66 | 3.47 |
|  |  | R2: | 107 ± 29.2 |  | -1.72 | -3.70 | -5.42 | 2.19 |
|  | *Mm* cGAS | R1: | 91.4 ± 58.9 | 104.5 ± 45.6 | -4.75 | -0.76 | -5.51 | 0.99 |
|  |  | R2: | 124 ± 72.0 |  | -5.18 | -0.25 | -5.33 | 1.05 |
| **Ap<sub>4</sub>A**<br>**vs.** | *Hs* cGAS | R1: | 126 ± 38.3 | 151.5 ± 28.7 | -4.34 | -0.98 | -5.32 | 1.28 |
|  |  | R2: | 184 ± 43.3 |  | -4.16 | -0.94 | -5.10 | 1.28 |
|  | *Mm* cGAS | R1: | 35.2 ± 9.2 | 41.4 ± 6.7 | -4.37 | -1.63 | -5.91 | 0.88 |
|  |  | R2: | 48.6 ± 9.9 |  | -4.08 | -1.81 | -5.89 | 1.16 |
| **Ap<sub>3</sub>A**<br>**vs.** | *Hs* cGAS | R1: | 92.3 ± 30.1 | 51.0 ± 6.4 | -1.24 | -4.27 | -5.51 | 1.57 |
|  |  | R2: | 49.1 ± 6.5 |  | -1.06 | -4.82 | -5.88 | 1.26 |
|  | *Mm* cGAS | R1: | 43.8 ± 5.8 | 41.5 ± 4.3 | -4.89 | -1.06 | -5.95 | 1.08 |
|  |  | R2: | 38.8 ± 6.3 |  | -5.35 | -0.67 | -6.02 | 1.03 |
| **Ap<sub>4</sub>G**<br>**vs.** | *Hs* cGAS | R1: | 111 ± 24.7 | 109.7 ± 14.9 | -8.11 | 2.71 | -5.40 | 1.41 |
|  |  | R2: | 109 ± 18.6 |  | -14.9 | 9.54 | -5.40 | 0.82 |
|  | *Mm* cGAS | R1: | 10.0 ± 0.4 | 10.06 ± 0.36 | -11.6 | 4.81 | -6.82 | 0.65 |
|  |  | R2: | 10.3 ± 0.8 |  | -12.1 | 5.33 | -6.80 | 0.63 |

Averaged $K_d$ ( $\mu\text{M}$ ) was calculated as weighted mean with propagated uncertainty.
R1 and R2 indicate independent replicates.

**Table S3.1.** LC-MS/MS acquisition parameters for targeted nucleotide and dinucleoside polyphosphate analysis. Mass transitions, collision energies, cell accelerator voltages, dwell times were optimized using chemically pure standards for each target analyte

| compound | precursor ion [m/z] | product ion [m/z] | dwell time [msec] | fragmentor voltage (V) | collision energy (V) | cell accelerator voltage (V) | polarity |  |
| --- | --- | --- | --- | --- | --- | --- | --- | --- |
| **Ap<sub>4</sub>G** | 851 | 504 | 40 | 380 | 33 | 5 | negative | qualifier |
| **Ap<sub>4</sub>G** | 851 | 487.9 | 40 | 380 | 34 | 5 | negative | quantifier |
| **Ap<sub>4</sub>A** | 835.1 | 488 | 40 | 380 | 31 | 5 | negative | qualifier |
| **Ap<sub>4</sub>A** | 835.1 | 407.8 | 40 | 380 | 38 | 5 | negative | quantifier |
| **Ap<sub>3</sub>A** | 755.1 | 407.8 | 40 | 380 | 34 | 5 | negative | qualifier |
| **Ap<sub>3</sub>A** | 755.1 | 272.8 | 40 | 380 | 47 | 5 | negative | quantifier |
| **GTP** | 522.1 | 424 | 40 | 380 | 22 | 5 | negative | qualifier |
| **GTP** | 522.1 | 158.9 | 40 | 380 | 41 | 5 | negative | quantifier |
| **ATP** | 505.9 | 407.9 | 40 | 380 | 21 | 5 | negative | qualifier |
| **ATP** | 505.9 | 158.8 | 40 | 380 | 28 | 5 | negative | quantifier |
| **GDP** | 442.1 | 344 | 40 | 380 | 19 | 5 | negative | qualifier |
| **GDP** | 442.1 | 159 | 40 | 380 | 29 | 5 | negative | quantifier |
| **ADP** | 425.9 | 327.8 | 40 | 380 | 17 | 5 | negative | qualifier |
| **ADP** | 425.9 | 133.9 | 40 | 380 | 22 | 5 | negative | quantifier |
| **GMP** | 362.1 | 211 | 40 | 380 | 14 | 5 | negative | qualifier |
| **GMP** | 362.1 | 79 | 40 | 380 | 39 | 5 | negative | quantifier |
| **AMP** | 346 | 96.9 | 40 | 380 | 24 | 5 | negative | qualifier |
| **AMP** | 346 | 78.9 | 40 | 380 | 30 | 5 | negative | quantifier |

